# ATG9 vesicles are a subtype of intracellular nanovesicle

**DOI:** 10.1101/2024.09.12.612637

**Authors:** Mary Fesenko, Daniel J. Moore, Peyton Ewbank, Elizabeth Courthold, Stephen J. Royle

## Abstract

Cells are filled with thousands of vesicles, which mediate protein transport and ensure homeostasis of the endomembrane system. Distinguishing these vesicles functionally and molecularly represents a major challenge. Intracellular nanovesicles (INVs) are a large class of transport vesicles that likely comprises of multiple subtypes. Here, we define the INV proteome and find that it is molecularly heterogeneous, and enriched for transmembrane cargo molecules including integrins, transporters, and ATG9A, a lipid scramblase associated with autophagy. ATG9A is known to reside in ‘ATG9 vesicles’: small vesicles that contribute to autophagosome formation. Using in-cell vesicle capture assays we found that ATG9A, as well as other ATG9 vesicle cargos, were in INVs. Quantitative analysis showed that virtually all ATG9 vesicles are INVs, but that only ∼20% of INVs are ATG9 vesicles, suggesting that ATG9 vesicles are in fact a subtype of INV, which we term ATG9A-flavor INVs. Finally, we show that perturbing ATG9A-flavor INVs impaired the autophagy response induced by starvation.

## Introduction

Membrane trafficking is a fundamental cellular process by which cargo molecules are moved between compartments via vesicular carriers. Several types of vesicle have been identified so far. Well-studied examples include those identified by their electron dense coat: clathrin-coated vesicles or COPII-coated vesicles (Boni-facino and Glick, 2004). However, the cell is filled with thousands of small, uncoated vesicles, whose roles in intracellular trafficking are poorly understood. We recently described a novel type of transport vesicle, intra-cellular nanovesicles (INVs). They are small (∼35 nm diameter), uncoated vesicles that are defined by the presence of Tumor Protein D52-like (TPD52-like) protein family members on their surface (Larocque et al., 2020, 2021; Larocque and Royle, 2022). INVs are an integral part of the intracellular trafficking network and they transport cargo on the anterograde and recycling pathways, mainly moving by diffusion (Larocque and Royle, 2022; Sittewelle and Royle, 2024). Importantly, INVs are likely a superfamily of vesicles with different identities, or flavors. For example, they collectively contain at least 16 different Rab GTPases and a minimum of four different R-SNAREs, which suggest the vesicles have diverse origins (Larocque et al., 2020). Disambiguation of the different flavors of INVs is therefore a major challenge.

There are four TPD52-like proteins (TPD52, TPD53/TPD52L1, TPD54/TPD52L2, TPD55/TPD52L3) that each contain a coiled-coil domain through which they can homo- or hetero-dimerize (Byrne et al., 1998), and four amphipathic helices, with a preference for high curvature membranes (Reynaud et al., 2022; Larocque et al., 2021). The third amphipathic helix in TPD54 conforms to an amphipathic lipid packing sensor (ALPS) motif, and mutation of a single positively charged residue in this region (R159E) prevents the otherwise tight association of TPD54 with INVs (Reynaud et al., 2022; Larocque et al., 2021). Due to its high expression among TPD52-like proteins, TPD54 is used as a marker for INVs (Larocque et al., 2020).

Autophagy, the cellular process responsible for clearing unnecessary or dysfunctional components, relies on the formation of autophagosomes. These specialized organelles encapsulate targeted cargoes and transport them to lysosomes for degradation. Of the core set of Atg proteins that are essential for autophagy, Atg9/ATG9A is the only transmembrane protein (Nishimura and Tooze, 2020), and functions as a lipid scramblase (Maeda et al., 2020; Guardia et al., 2020; Matoba et al., 2020). In mammalian cells, ATG9A resides on small uncoated vesicles, termed ‘ATG9 vesicles’, that cycle between the *trans*-Golgi network, the plasma membrane and the endosomal system (Orsi et al., 2012; Popovic and Dikic, 2014; Young et al., 2006). During autophagy, ATG9 vesicles move to the phagophore assembly site, where they contribute to phagophore formation through mechanisms that are still being determined (Orsi et al., 2012; Karanasios et al., 2016; Olivas et al., 2023; Broadbent et al., 2023). ATG9 vesicles are formed at the *trans*-Golgi network (TGN) via the recruitment of adaptor proteins, of which the AP-4 dependent mechanism is the best characterized (Davies et al., 2018; Mattera et al., 2017). Neurological diseases associated with AP-4 deficiency, might therefore be explained by dysregulation of autophagy caused by mistrafficking of ATG9A. Finally, ATG9 vesicles may have additional, autophagy-independent functions. For example, in plasma membrane repair (Claude-Taupin et al., 2021), in mobilization of lipids from lipid droplets to mitochondria (Mailler et al., 2021), and in integrin trafficking during cell migration (Campisi et al., 2022). At least superficially, ATG9 vesicles resemble INVs: their size, lack of coat, their diffusive movement through the cytoplasm and their involvement in exocytosis and integrin recycling (Broadbent et al., 2023; Holzer et al., 2024; Sittewelle and Royle, 2024; Larocque et al., 2021).

To determine the different molecular identities of INV and how these subtypes might relate to other vesicle types, we set out to determine the INV proteome. Analysis of the proteome underscored the molecular heterogeneity of this class of transport vesicle. The presence of ATG9A and other ATG9 vesicle cargos in the proteome prompted us to investigate whether ATG9 vesicles could be one flavor of INV. We show that a fraction of INVs can account for the majority of ATG9 vesicles and that perturbing INVs results in impaired autophagy.

## Results

### Determining the INV proteome

Our previous work suggested that INVs could be immunoisolated from cells and analyzed proteomically (Larocque et al., 2020, 2021). We therefore optimized an INV immunoisolation procedure from cells permeabilized with low concentrations of digitonin, and using a GFP nanobody (GFP-Trap) to isolate the INVs by virtue of the marker protein GFP-TPD54. For optimization, three different cell backgrounds were used, either parental HeLa cells (control), HeLa cells stably expressing GFP-TPD54 (WT) or a GFP-TPD54 mutant (R159E) that cannot bind INVs (Figure 1A-D). The immunoisolated material from GFP-TPD54 WT cells was significantly enriched for 111 or 120 proteins when compared with control or cells expressing GFP-TPD54 R159E, respectively (Figure 1B,D). The enriched proteins included all TPD52-like proteins expressed in HeLa cells (TPD54, TPD52, and TPD53) as well as various Rabs, VAMPs and cargo proteins, suggesting successful purification of INVs. By contrast, immunoisolated material from cells expressing GFP-TPD54 R159E did not isolate INVs, with only 17 proteins alongside TPD54 significantly enriched compared to control (Figure 1C and S1A,B). Nine of these proteins were known contaminants in affinity purification experiments (Mellacheruvu et al., 2013). This suggests that the majority of proteins enriched following immunoisolation from GFP-TPD54 WT expressing cells are *bona fide* INV proteins and are not, for example, binding directly to TPD54.

**Figure 1.**
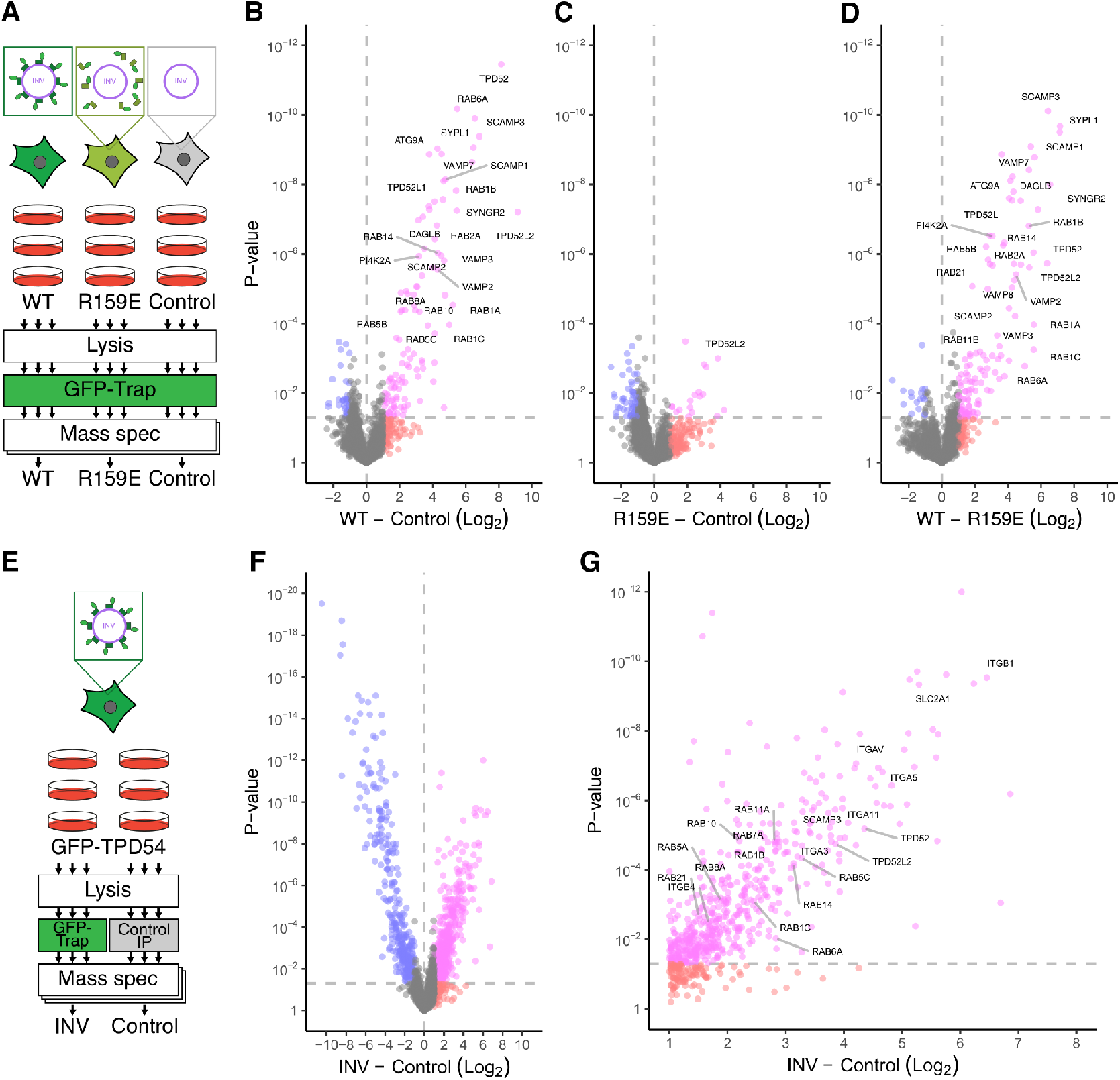
Determining the INV proteome. (**A**) Schematic diagram of INV immunoisolation: GFP-Trap pulldown from HeLa cells stably expressing GFP-TPD54 WT, GFP-TPD54 R159E, or untransfected Hela cells (Control). (**B-D**) Volcano plots to show comparisons of (B) WT and Control, (C) R159E and Control, or (D) WT and R159E. Data are from two independent runs consisting of three replicates each. (**E**) Schematic diagram of INV immunoisolation: GFP-Trap or control IP pulldown from GFP-TPD54 knock-in HeLa cells. (**F**) Volcano plot to show the comparison of INV and Control isolation. (**G**) Expanded view of F. Data are from three independent runs of three replicates each. Colors indicate: pink, fold-change > 2 and p < 0.05; red, fold-change > 2 and p > 0.05; blue, fold-change < 2 and p < 0.05; gray, remainder.

With this method in hand, we extended the approach to large-scale isolation of INVs from a knock-in GFPTPD54 HeLa cell line. This allowed us to isolate endogenous INVs via GFP-Trap and compare to a control IP from a single cell line source (Figure 1E). Analysis of these experiments yielded 525 proteins that were significantly enriched compared to the control (Figure 1F-G). Again, TPD52-like proteins, Rabs and cargo were present alongside other proteins indicating deeper coverage with this strategy. We combined the data from both approaches to obtain a list of 602 proteins that we consider a first INV proteome (Supplementary Table S1).

## Exploring the INV proteome

To begin exploring the INV proteome, we used PANTHER protein classification to categorize INV proteins (Figure 2A and Supplementary Table S2). The “membrane traffic protein” class and “G-protein” subclass were both large with 35 proteins each. The INV proteome contains 17 Rab GTPases (Rab1A-C, Rab2A, Rab5A-C, Rab6A, Rab7A, Rab8A-B, Rab10, Rab11A-B, Rab14, Rab21, and Rab34), 4 VAMPs (VAMP2, VAMP3, VAMP7, and VAMP8) as well as TPD52-like proteins; which is consistent with previous vesicle trapping experiments (Larocque et al., 2020). Aside from dynamin-2 (DNM2) and coatomer subunit-beta (COPB1), there was an absence of machinery associated with classical coated vesicles. The largest category was without PANTHER protein classification (Unclassified) and this contained many secreted proteins (see below). When used as bait in proximity biotinylation proteomics studies, a number of INV proteins, including DNAJC5, LAMTOR1, LAMP2, RhoB, Rab2, Rab5C, Rab11A, and NRAS, have each identified TPD54 as their enriched putative interactor (Barker et al., 2024; Wilson et al., 2023; Go et al., 2021; Adhikari and Counter, 2018), giving confidence to our dataset. Cargo proteins dominate the proteome. Across multiple categories, receptors, transporters and other trans-membrane cargo proteins feature strongly. Examples include the transferrin receptor (TFRC), mannose-6-phosphate receptors (M6PR, IGF2R), and integrins (ITGA3, ITGA5, ITGA6, ITGAV, ITGA11, ITGB1) some of which have previously been shown to be trafficked via INVs (Larocque et al., 2020, 2021).

**Figure 2.**
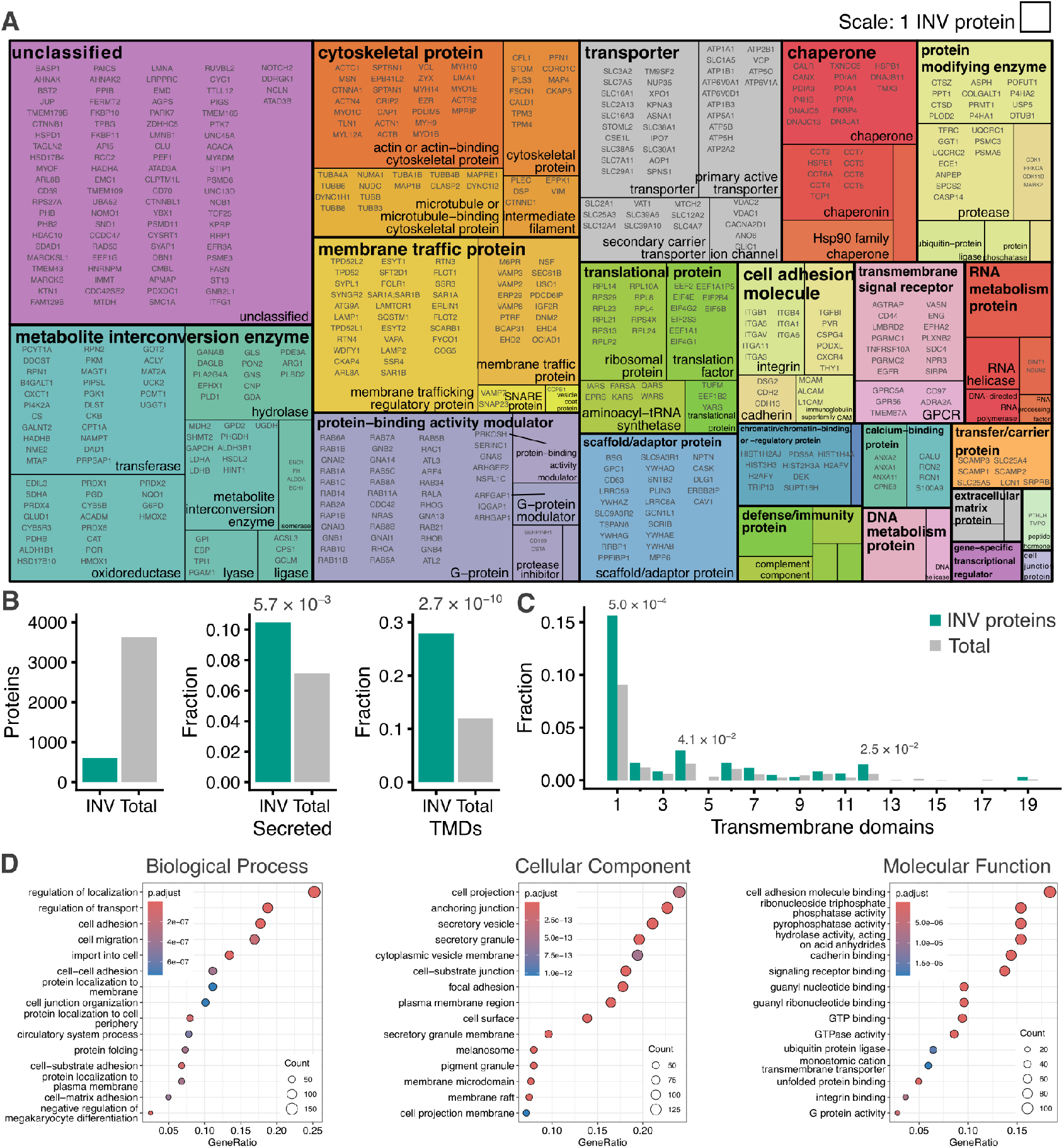
Composition of the INV proteome. (**A**) Treemap visualization of classification of 602 INV proteins. Two levels of PANTHER classification are shown (for details see Materials and Methods, for full list see Supplementary Table S2). (**B**) Bar charts showing the INV proteins versus all proteins detected (Total), and the fractions of INV or total proteins, which are designated as secreted or which contained one or more transmembrane domain (TMD). Enrichment of secreted or TMD-containing proteins in INVs versus total is indicated, p-values from Pearson’s Chi-squared test. (**C**) Fractions of proteins with the indicated number of transmembrane domains. Significant enrichment is indicated by p-values from Pearson’s Chi-squared test. (**D**) Gene Ontology (GO) term enrichment analysis for 602 INV proteins versus total.

We found that the INV proteome had a significant enrichment of secreted proteins (10.5% vs 7.2%, χ^2^ = 7.66, p = 5.7 × 10^−3^) and of proteins with one or more transmembrane (TM) domains (28.0% vs 17.0%, χ^2^ = 39.9, p = 2.7 × 10^−10^), which is consistent with their previously characterized role as transport vesicles on the anterograde and recycling pathways (Figure 2B). Of the INV proteins that have TM domains, there was a notable enrichment of proteins with either 1, 4, or 12 TM domains (Figure 2C). The single pass protein group was the largest and was heterogeneous. However, the 12 TM domain proteins were all transporters (SLC12A2, SLC12A4, SLC16A1, SLC16A3, SLC2A1, SLC2A13, SLC7A11, SLC7A5, SPNS1), while the 4 TM domain proteins include tetraspanins (CD63, TSPAN6), and the secretory carrier-associated membrane proteins (SCAMP1, SCAMP2, SCAMP3), synaptogyrin-2 (SYNGR2), and synaptophysin-like protein 1 (SYPL1) whose neuronal counterparts are usually associated with synaptic vesicles. Gene Ontology (GO) enrichment analysis highlighted the cellular components secretory vesicle and secretory granule among other terms including cell migration, cell adhesion, and GTPase binding (Figure 2D).

We next compared the INV proteome with previously published vesicle proteomic datasets. The presence of VAMP2, SCAMPs, SYNGR2 and SYPL-1, together with vacuolar-type ATPase subunits (ATP6V0A1, ATP6V0D1) was intriguing, especially since TPD52-like proteins are abundant in published synaptic vesicle proteomes (Bradberry et al., 2022; Biesemann et al., 2014). This likely reflects the fraction of INVs that are exocytosed, as we previously demonstrated (Sittewelle and Royle, 2024). First, given the non-neuronal origin of our INV proteome we compared our data with that from synaptic-like microvesicles from PC12 cells, rather than synaptic vesicles (Salazar et al., 2005). We found significant enrichment in the INV proteome for proteins from this dataset (5.5% vs 1.8%, χ^2^ = 28.44, p = 9.653 × 10^−8^); with 27.5% of SLMV proteins featured in the INV proteome (Supplementary Figure S2A, D).

Since the lipid scramblase ATG9A was also among the cargo proteins in the INV proteome we next compared our data to an ATG9 vesicle proteome (Judith et al., 2019). We found a significant enrichment in the INV proteome for proteins in the ATG9 vesicle dataset (23.8% vs 10.7%, χ^2^ = 78.73, p < 2.2 × 10^−16^). Moreover, the intensity or enrichment of common proteins from the two datasets were correlated (ρ = 0.21, p < 0.011, Supplementary Figure S2B, D). Interestingly, in the ATG9 dataset, the INV marker protein TPD54 was one of the most enriched proteins in conditions that upregulate autophagy (Judith et al., 2019), yet the majority of the proteins common to the two datasets were de-enriched upon starvation (Supplementary Figure S2C). Finally, as a control, we tested for enrichment in the INV proteome for proteins in a CCV proteomic dataset (Borner et al., 2012) and found no enrichment (1.5% vs 1.7%, χ^2^ = 0.02, p = 0.89, Supplementary Figure S2D).

In summary, this analysis strengthens the view that INVs are a mix of different subtypes. The enrichment of ATG9 vesicle proteins in the INV proteome and the differential response of the INV proteome to starvation indicated to us that ATG9 vesicles could be a subtype of INV.

### ATG9A is in intracellular nanovesicles

To investigate whether ATG9A is in INVs, we used a vesicle capture imaging assay. Here, INVs are captured at the mitochondria using induced heterodimerization between the FKBP-tagged INV marker protein (GFP-FKBP-TPD54) and MitoTrap (Mito-mCherry-FRB[T2098L]) with rapalog (AP21967, 5 µM). The co-relocation of another protein coincident with TPD54 indicates it is present in INVs (Figure 3A). Detection of endogenous ATG9A by immunofluorescence showed that it was strongly co-relocated to the mitochondria when GFP-FKBP-TPD54 WT was relocalized there; but not when GFP-FKBP or GFP-FKBP-TPD54 R159E mutant were relocalized (Figure 3B). Quantification of the ratio of mitochondrial fluorescence versus the total fluorescence per cell demonstrated successful relocalization of the GFP-FKBP tagged proteins to the mitochondria in each case and underscored that co-relocation of ATG9A only occurred with TPD54 WT (Figure 3C). These results indicate that endogenous ATG9A is found in INVs in cells.

**Figure 3.**
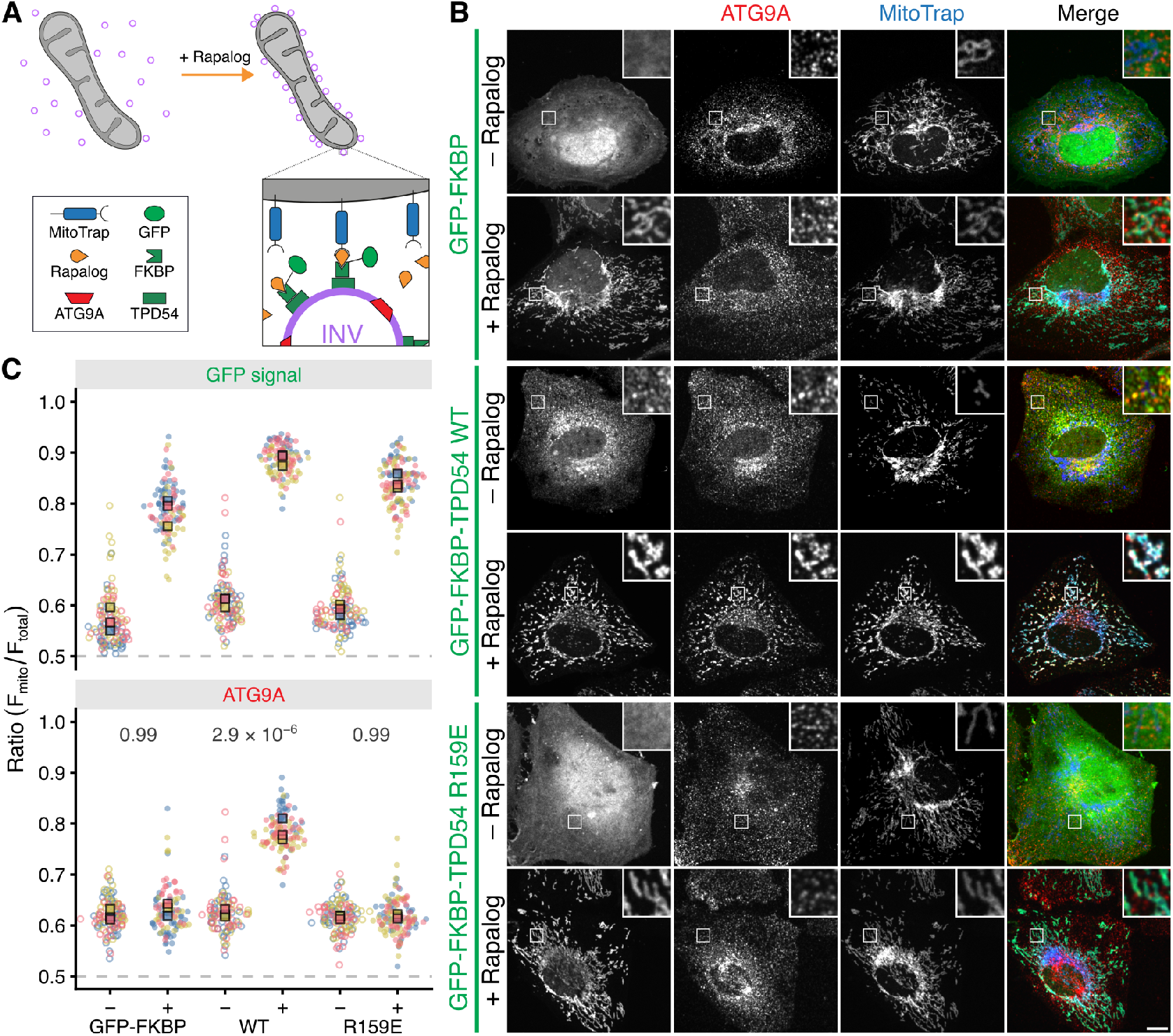
ATG9A is in intracellular nanovesicles. (**A**) Schematic diagram of vesicle capture at mitochondria as a test for co-relocation. MitoTrap is an FRB domain targeted to mitochondria, GFP-FKBP-TPD54 is coexpressed and, when rapalog is added, the INVs associated with TPD54 become trapped at the mitochondria. Any protein that is also in the INVs is co-relocated with TPD54 (Larocque et al., 2020). (**B**) Representative confocal images of HeLa cells expressing MitoTrap (pMito-mCherry-FRB T2098L, blue) and either GFP-FKBP, GFP-FKBP-TPD54 WT, or GFP-FKBP-TPD54 R159E (green), stained with AF647-conjugated anti-ATG9A antibody (red). Cells were treated with rapalog (5 µM, 5 min) or not, as indicated. Scale bar, 10 µm; insets, 4 *×* zoom. (**C**) Superplot to show the relocalization of GFP-FKBP or GFP-FKBP-tagged TPD54 construct and the extent of co-relocation of ATG9A. Data are expressed as a fraction of the total fluorescence per cell that is at the mitochondria. Spots indicate individual cell measurements (n = *∼*50 per repeat), colors indicate independent experimental repeats (n = 3), squares show the mean value for each replicate. P-values, two-way ANOVA with Tukey’s HSD post-hoc test.

### ATG9 vesicles are INVs

Next, we performed the reciprocal test: relocalizing ATG9A and asking if TPD54 is co-relocated. To do this, ATG9A-containing vesicles were captured at the mitochondria using ATG9A-FKBP-mCherry with Mito-Trap (Mito-EBFP2-FRB T2098L). Using this approach we found that GFP-TPD54 WT but not the R159E mutant was co-relocated to the mitochondria, when ATG9A-FKBP-mCherry was relocalized (Figure 4A,B). This result indicates that GFP-TPD54 WT is present on the same vesicle as ATG9A-FKBP-mCherry by virtue of membrane-binding rather than by an association with ATG9A itself. Moreover, endogenous TPD54 (in GFP-TPD54 knock-in cells) was also co-relocated with ATG9A-FKBP-mCherry, indicating that co-relocation was not a result of overexpression of GFP-TPD54 (Figure 4C). Importantly, the co-relocation of TPD54 was dependent on ATG9A-containing vesicle capture, because relocalization of FKBP-mCherry had no effect on GFP-TPD54 localization (Figure 4A,C). Quantification of the ratio of mitochondrial TPD54 fluorescence postversus pre-rapalog treatment confirmed the specific co-relocation of TPD54 with ATG9A relocalization (Figure 4D). The results indicate that relocalization–corelocation of ATG9A and TPD54 was reciprocal.

**Figure 4.**
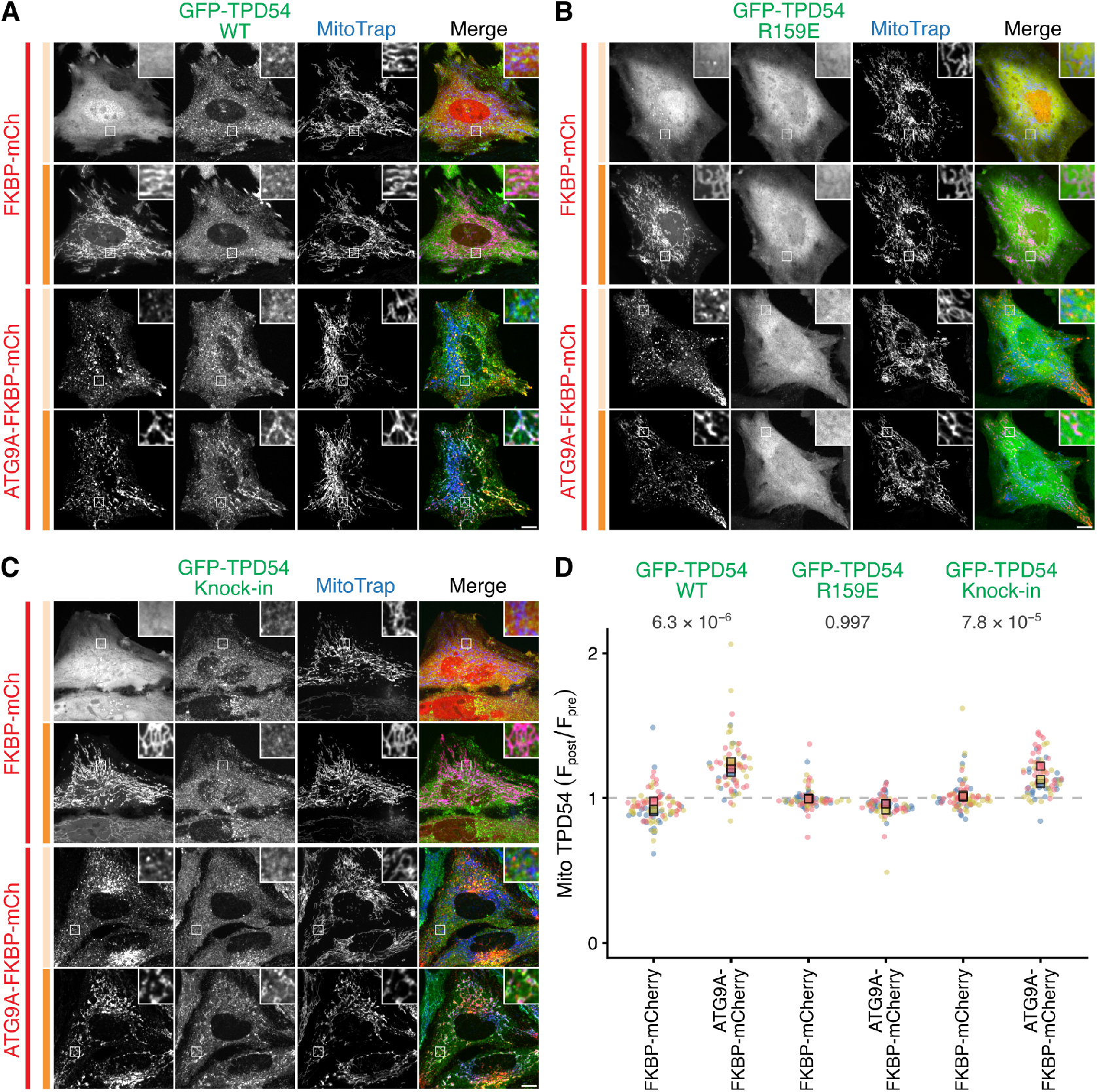
ATG9 vesicles have the INV marker TPD54. (**A-C**) Representative confocal images of HeLa cells expressing MitoTrap (pMito-EBFP2-FRB T2098L, blue) and either FKBP-mCherry or ATG9A-FKBP-mCherry (red). Cells are co-expressing GFP-TPD54 WT (A), GFP-TPD54 R159E (B), or endogenous GFP-TPD54 knock-in (C) (green). For each condition, the same cell is shown pre(pale orange bar) and post-treatment (dark orange bar) with rapalog (5 µM, 5 min). Scale bar, 10 µm; insets, 4 *×* zoom. (**D**) Superplot to show the ratio of mitochondrial fluorescence of GFP-TPD54 post- vs pre-rapalog treatment. Spots indicate individual cell measurements (n = 21 - 69 per repeat), colors indicate independent experimental repeats (n = 3), squares show the mean value for each replicate. P-values, two-way ANOVA with Tukey’s HSD post-hoc test. Confirmation of FKBP-mCherry and ATG9A-FKBP-mCherry relocalization is shown in Supplementary Figure S3.

The most likely interpretation of these experiments is that the two proteins are present in the same vesicles. Since our analysis was done at two discrete time points, it is formally possible that distinct INVs were captured secondarily to the trapping of ATG9 vesicles at the mitochondria. To address this point, we used live-cell imaging to study the kinetics of ATG9A-FKBP-mCherry relocalization and endogenous GFP-TPD54 co-relocation to the mitochondria in response to 5 µM rapalog (Figure 5A-D, Supplementary Movie SV1). Analysis of the mitochondrial fluorescence of each protein as a ratio of the pre-rapalog fluorescence, revealed a jump in fluorescence which was coincident in both channels (Figure 5A-C). The kinetics of relocalization and co-relocation were highly similar: the time-constant (*τ*) of a single exponential function fitted to each protein co-varied across 34 cells from three experiments (*R*^2^ = 0.774, Figure 5D). This strengthens the evidence that the two proteins are on the same vesicles and argues against the secondary capture of a distinct vesicle type. This analysis also revealed a disparity in the extent of co-relocation of TPD54 compared to the amount of ATG9A that was relocalized (see below). Together these results indicate that TPD54 is found in ATG9A-containing vesicles in cells. Since our definition of INVs is that they contain TPD54, then it follows that ATG9 vesicles are actually INVs.

**Figure 5.**
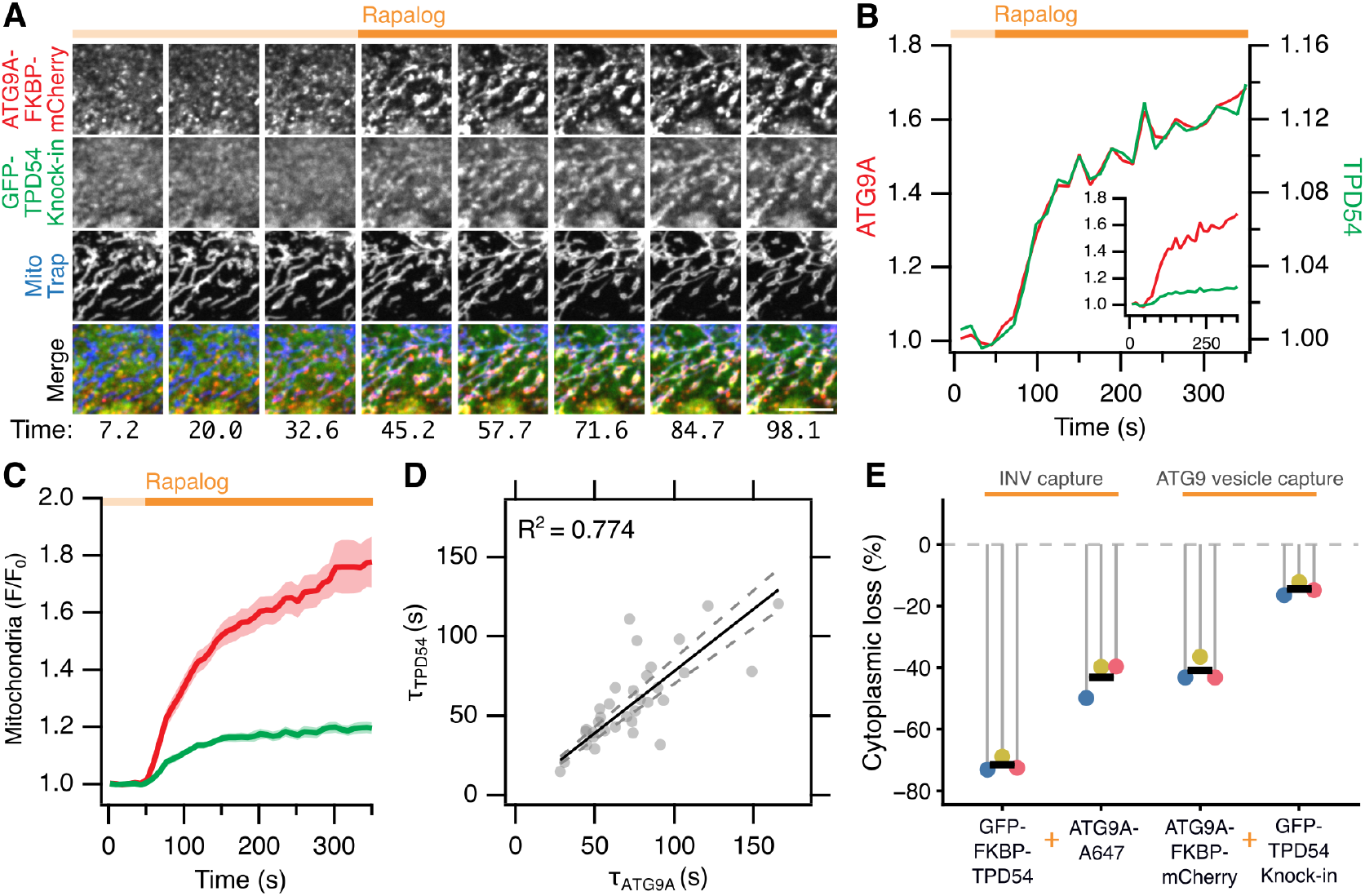
ATG9 vesicles are a subset of INVs. (**A**) Cropped stills from a movie of GFP-TPD54 (green) knock-in cells expressing ATG9A-FKBP-mCherry (red) and MitoTrap (blue); capture of ATG9A-positive vesicles at the mitochondria is induced by rapalog (5 µM at 40 s). Scale bar, 5 µm. See Supplementary Video SV1. (**B**) Quantification of mitochondria fluorescence of ATG9A-FKBP-mCherry (left axis) and GFP-TPD54 (right axis) for the cell shown in A. Plot shows fluorescence divided by initial fluorescence (F/F0) Inset: traces shown on same scale. (**C**) Average mitochondrial ATG9A-FKBP-mCherry and GFP-TPD54 signal for 34 cells, from 3 independent experiments (mean ±sem). (**D**) Plot of the time constant (Tau) for single exponential fits to the ATG9A-FKBP-mCherry and the GFP-TPD54 traces. Lines show a linear fit to the data and 95% confidence bands; *R*^2^ value for the fit through the origin is shown. (**E**) Loss of cytoplasmic signal following relocalization. Lollipops show the average decrease per experimental repeat (n = 31 - 52 cells), bar shows the mean (n = 3 independent repeats). Data are calculated from experiments in Figures 3 and 4.

### ATG9A-containing vesicles are subset of INVs

In the relocalization–co-relocation experiments, the loss of protein from the cytoplasm can be quantified in order to estimate the sizes of the vesicle pools that are captured at the mitochondria. We therefore compared the loss of fluorescence in the cytoplasm after adding rapalog as a percentage of its steady-state localization. During INV capture – when TPD54 was relocalized and ATG9A co-relocated – there was a 71.5% decrease in cytosoplasmic TPD54 resulting in a 43.1% decrease in endogenous ATG9A signal (Figure 5E). During capture of vesicles containing ATG9A – when ATG9A was relocalized and TPD54 co-relocated – we found a 41.0% decrease in cytoplasmic ATG9A causing only a 14.5% decrease of the endogenous TPD54. This means that the size of the pool of vesicles that can be relocalized is ∼70% for TPD54 and ∼40% for ATG9A. Therefore, the capture of INVs results in trapping of the entire ATG9 vesicle pool; whereas only ∼20% of the INV pool was trapped when all of the ATG9A-positive vesicles were captured. That is, all ATG9 vesicles are INVs, but not all INVs are ATG9 vesicles. We conclude that ATG9 vesicles are a subset of INVs and we term them ATG9Aflavor INVs.

### ATG9 vesicle cargo is also in INVs

If ATG9 vesicles are ATG9A-flavor INVs, then we would predict that other ATG9 vesicle cargos would also be found in INVs. To test this possibility, we examined the co-relocation of five candidate proteins reported to be present on ATG9 vesicles: SH3GLB1, ARFIP2, DAGLB, PI4K2A, PI4KB (Takahashi et al., 2011; Judith et al., 2019; Davies et al., 2022). Of these, DAGLB and PI4K2A were enriched in the INV proteome (Supplementary Table S1). We performed vesicle capture using relocalization of either ATG9A-FKBP-mCherry or mCherry-FKBP-TPD54 to MitoTrap and used FKBP-mCherry or mCherry-FKBP-TPD54 R159E mutant as the respective negative controls for the specificity of cargo co-relocation due to vesicle capture. For GFP-tagged SH3GLB1, DAGLB and PI4K2A, we observed robust co-relocation that was caused by the relocalization of ATG9A or of TPD54 (Figure 6A,B). No co-relocation was observed for GFP-tagged ARFIP2 nor for PI4KB when either ATG9A-FKBP-mCherry or mCherry-FKBP-TPD54 was relocalized to mitochondria (Figure 6B). This was unexpected because ARFIP2 and PI4KB strongly colocalized with ATG9A at the TGN prior to re-localization (Supplementary Figure S4). This might indicate that these two proteins are associated with ATG9A at the TGN and that this pool of ATG9A is not relocalized in our experiments, or they are not stably associated with vesicles, such that they fall off during vesicle trapping. Where co-relocation of cargo proteins was observed, it was specific because relocalization of FKBP-mCherry or mCherry-FKBP-TPD54 R159E had no effect any of the candidates. The finding that the same candidate proteins were co-relocated by relocalization of ATG9A or of TPD54 underscores the idea ATG9 vesicles are ATG9A-flavor INVs. The magnitude of co-relocation of SH3GLB1, DAGLB and PI4K2A was greater with relocalization of TPD54 than with ATG9A, which indicates that these proteins may be on other INV subtypes in addition to ATG9A-flavor INVs.

**Figure 6.**
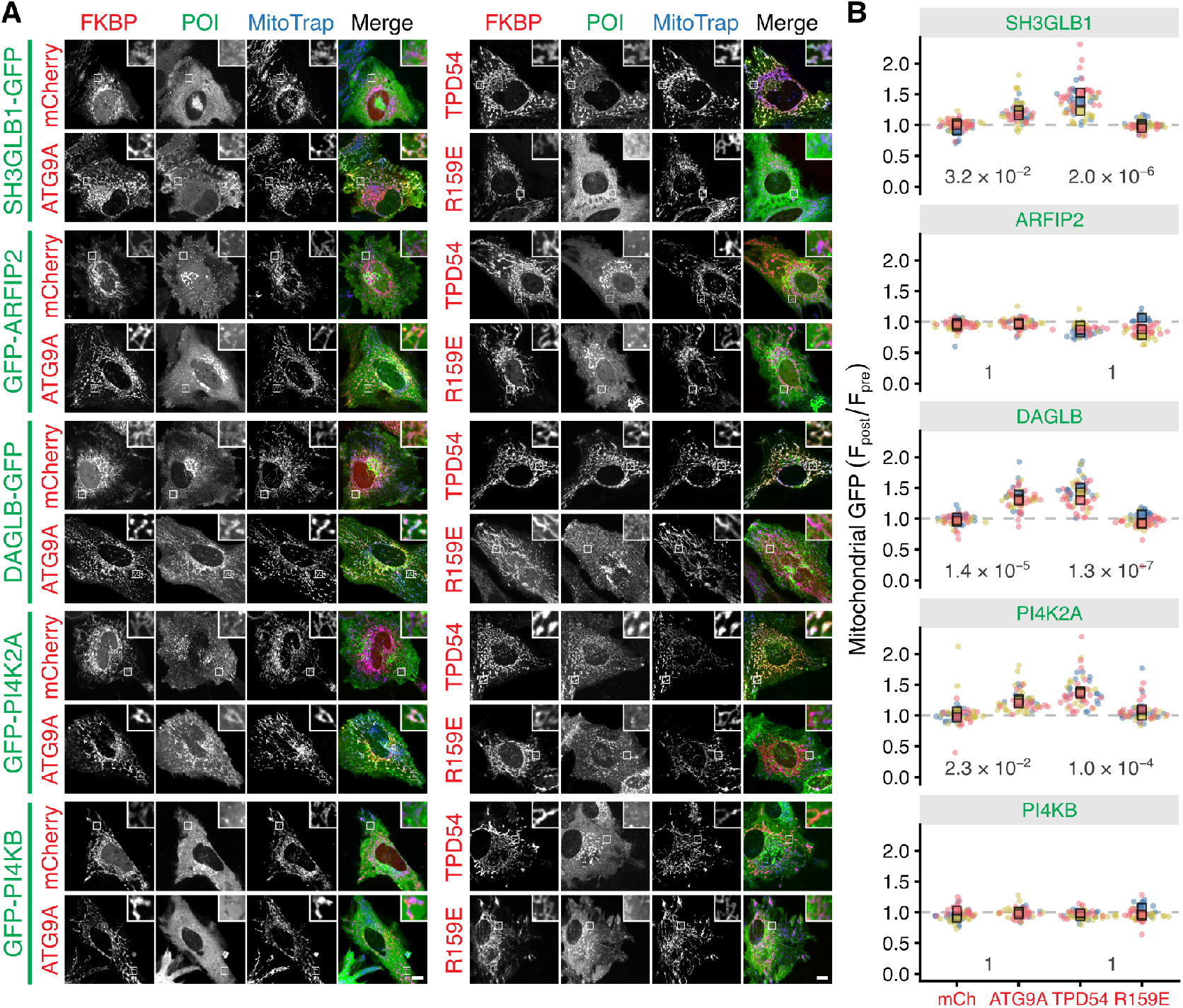
ATG9A-flavor INVs have ATG9 vesicle cargos. (**A**) Representative confocal images of HeLa cells expressing MitoTrap (pMito-EBFP2-FRB T2098L, blue) and either FKBP-mCherry, ATG9A-FKBP-mCherry, mCherryFKBP-TPD54 WT, or mCherry-FKBP-TPD54 R159E (red). Cells are co-expressing GFP-tagged SH3GLB1, ARFIP2, DAGLB, PI4K2A, or PI4KB as indicated (green). For each condition, only the post-rapalog (5 µM, 5 min) treatment images are shown, for the pre-treatment images see Supplementary Figure S4. Scale bar, 10 µm; insets, 4 *×* zoom. (**B**) Superplots to show the ratio of mitochondrial fluorescence of the indicated GFP-tagged protein post- vs pre-rapalog treatment. Spots indicate individual cell measurements (n = 10 - 28 cells per repeat), colors indicate independent experimental repeats (n = 3), squares show the mean value for each replicate.

### ATG9A-flavor INVs can be distinguished from another INV subtype by selective transport

RUSC2 overexpression induces a striking aggregation of AP4-derived vesicles and their cargoes, including ATG9A, at the edges of the cell (Davies et al., 2018; Guardia et al., 2021). To test if ATG9A-flavor INVs respond to HA-RUSC2 overexpression in the same way, we examined vesicle distributions in GFP-TPD54 knock-in HeLa cells in these conditions by microscopy. Large accumulations of HA-RUSC2 at the cell periphery were found in the majority of cells, and these accumulations were positive for GFP-TPD54 and ATG9A immunofluorescence (Supplementary Figure S5. Another INV cargo – CIMPR/IGF2R – did not accumulate at HA-RUSC2/TPD54 vesicle aggregations (Supplementary Figure S5. CIMPR is found in our INV proteome and we have previously shown that CIMPR-containing INVs can be relocalized to the mitochondria by INV trapping (Larocque et al., 2020). Therefore, these results suggest two things. First, that endogenous ATG9Aflavor INVs behave in the same way as ATG9 vesicles; second, the selectivity of this manipulation highlights that not all INVs are ATG9A-flavor INVs, and that other subtypes of INV behave differently.

### Depletion of TPD54 impairs the autophagy response

Having established that ATG9 vesicles are a subset of INVs, we next tested whether these INVs were functional during autophagy. To do this, we depleted TPD54 using RNAi and examined the levels of the autophagy marker protein LC3B/MAP1LC3B in HeLa cells under fed or starvation conditions, with or without bafilomycin A1 (BafA1, 100 nM). Western blotting revealed that in control RNAi cells, the level of LC3B-II – the lipidated form of LC3B – was low in basal conditions (fed, no BafA1) and increased with BafA1 treatment, indicating a functional basal autophagic flux response (Figure 7A,B). When autophagy was upregulated by amino acid and growth factor starvation, the amount of LC3B-II increased by 2.8-fold, as expected. The level of LC3B-II increased further in cells that were starved and treated with BafA1, indicating the expected higher autophagic flux in response to starvation-induced autophagy unpregulation. In TPD54-depleted cells, the level of LC3B-II was higher in basal conditions, but the rate of basal autophagic flux was unchanged when compared to control RNAi cells. However, we did not observe further increases in the levels of LC3B-II in response to starvation in TPD54-depleted cells, and induced autophagic flux was also dampened, relative to that in control RNAi cells (Figure 7A,B).

**Figure 7.**
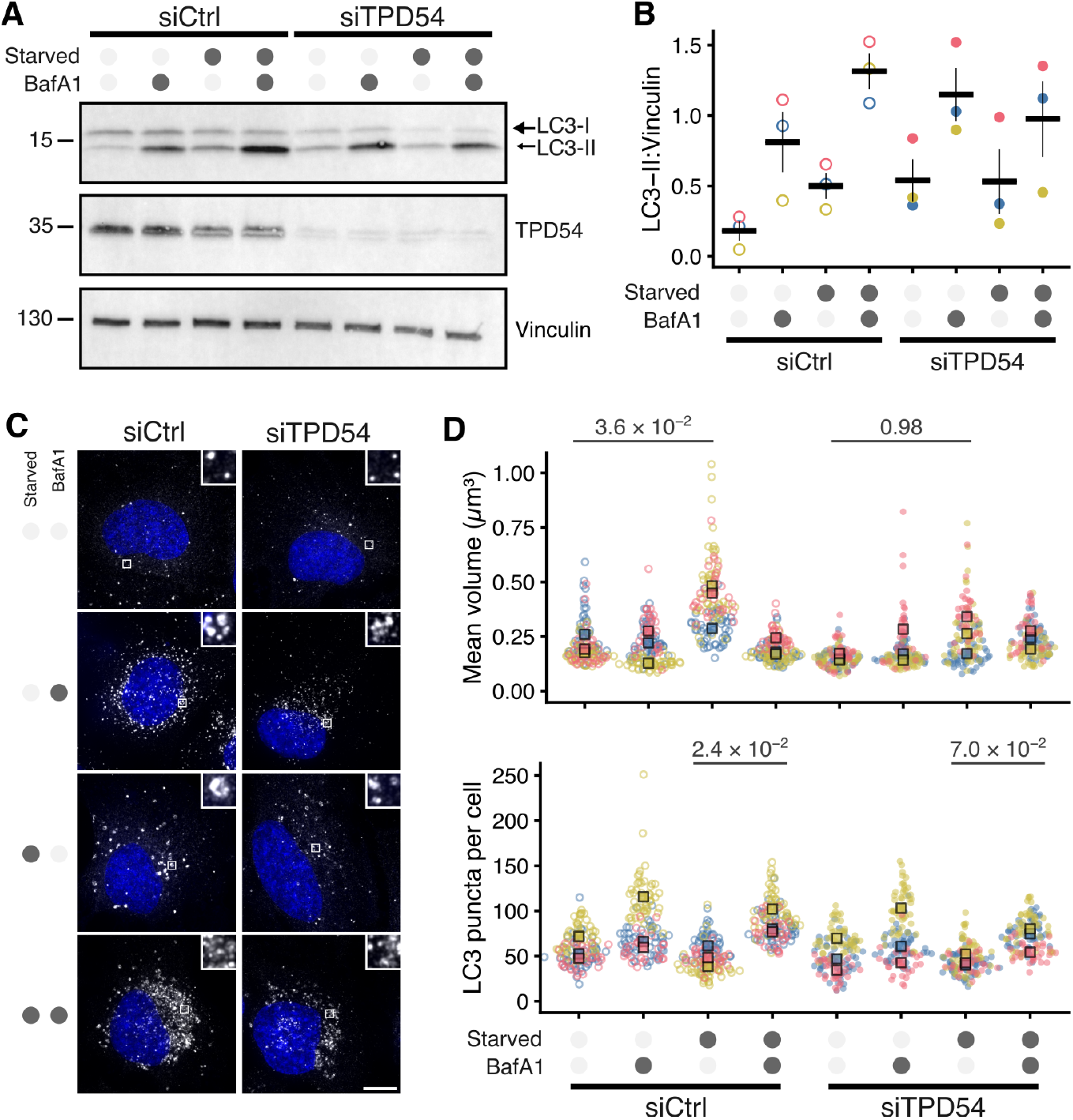
Depletion of TPD54 impairs the autophagy response. HeLa cells were transfected with control (GL2, siCtrl) or TPD54 (siTPD54) siRNA, then either starved (3 h) or not and either treated with BafA1 (100 µM) or not, as indicated. (**A**) Western blot showing levels of LC3B, TPD54 and vinculin (loading control). Result is typical of three repeats shown in Supplementary Figure S6. (**B**) Quantification of LC3B-II signal normalized to vinculin. Dots show experimental repeats and bars shown mean ± sem. (**C**) Representative maximum intensity z-projections showing LC3 immunofluorescence (white) and DAPI (blue). Scale bar, 10 µm; insets, 5 *×* zoom. TPD54 depletion for these experiments was confirmed by Western blot (Supplementary Figure S7). (**D**) Superplots to show the average volume and number of LC3 puncta. Spots indicate individual cell measurements (n = 30 - 50 cells per repeat), colors indicate independent experimental repeats (n = 3), squares show the mean value for each replicate. P-values from Tukey’s HSD post hoc test, following two-way ANOVA.

Under the same conditions, we also directly visualized autophagosome formation using immunofluorescence detection of LC3B. Starvation resulted in an increase in the mean volume of LC3 puncta per cell, but this increase was less pronounced in TPD54-depleted cells (Figure 7C,D). The mean volume of LC3 puncta in starved cells was 24.0–40.2% lower in TPD54-depleted cells than controls, indicating inhibition of autophagosome formation (Figure 7D). Basal autophagic flux, assessed by quantifying numbers of LC3 puncta per cell was unaffected by TPD54 RNAi, however autophagic flux induced by starvation was lower in TPD54 depleted cells. BafA1 treatment of starved, control RNAi cells resulted in a 0.3-to 1.6-fold accumulation of smaller LC3 puncta. In TPD54-depleted cells, the LC3 puncta accumulation in starved, BafA1-treated cells was less pronounced and the mean number of LC3 puncta per cell was 6.8–29.3% lower than in control RNAi cells (Figure 7D). Together, these data indicate that ATG9Aflavor INVs are functionally equivalent to previously described ATG9 vesicles during amino acid and growth factor starvation-induced autophagy. Perturbing INV function through TPD54 depletion inhibited autophagosome formation and impedes autophagic flux, identifying TPD54 as a novel molecular regulator of autophagy.

### Cellular origin of ATG9A-flavor INVs

ATG9A has been shown to be redistributed from the TGN upon starvation (Orsi et al., 2012). We therefore tested whether INVs were involved in this phenomenon by examining the effect of TPD54 depletion on starvation-induced ATG9A redistribution. Using immunofluorescence of ATG9A and TGN46, we saw that under fed conditions, a fraction of ATG9A overlaps with the TGN (Figure 8A). Following 3 h amino acid and growth factor starvation, the ATG9A signal at the TGN was significantly lower in control RNAi conditions, confirming the redistribution phenomenon (Figure 8A,B). However in TPD54-depleted cells, this redistribution was blocked and the levels of ATG9A at the TGN in starved cells were not significantly different to those in fed conditions (Figure 8A,B). If the redistribution of ATG9A from the TGN during starvation represents the exit ATG9A in new ATG9A-flavor INVs to mediate autophagosome formation, and this is blocked by TPD54 depletion, then it suggests that TPD54 is involved in their formation.

**Figure 8.**
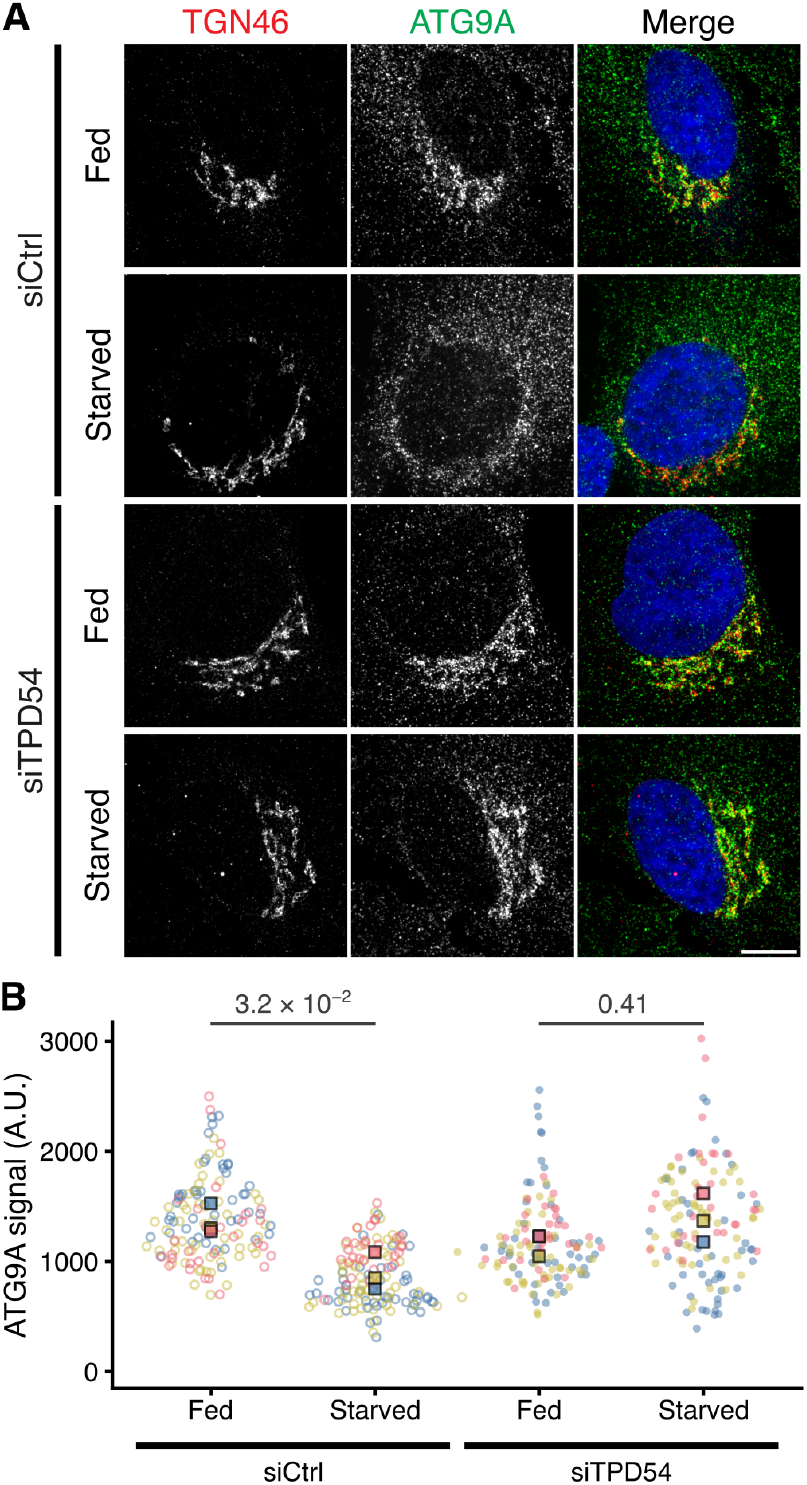
Starvation-induced loss of ATG9A from the TGN is blocked by TPD54 depletion. (**A**) Representative confocal micrographs of HeLa cells transfected with control (GL2, siCtrl) or TPD54-targeting siRNA (siTPD54) in either fed or starved (3 h) conditions as indicated, stained for TGN46 (red) and ATG9A (green). Scale bar, 10 µm. TPD54 depletion was confirmed by Western blot (Supplementary Figure S7). (**B**) Superplot to show the ATG9A immunofluorescence signal at the Golgi. Spots indicate individual cell measurements (n = 30 - 50 per repeat), colors indicate independent experimental repeats (n = 3), squares show the mean value for each replicate. P-values from Tukey’s HSD post hoc test, following two-way ANOVA.

## Discussion

Understanding the functions and identities of the thousands of uncoated vesicles inside cells is a major challenge. Here, we described the proteome of intracellular nanovesicles and found that this is a large, molecularly diverse class of vesicles, likely comprising of multiple INV subtypes. We showed that ATG9 vesicles – a key membrane source during autophagy – are a subtype of INV, and confirmed that it is these ATG9A-flavor INVs that are important for autophagic flux.

ATG9 vesicles have been classically defined by the presence of the transmembrane Atg gene product, ATG9A or ATG9B in humans. Our finding, that these vesicles are actually a subtype of INV, is based on multiple lines of evidence. First, ATG9A and other ATG9 vesicle cargos are found in the INV proteome, and the INV marker, TPD54, is found in an ATG9 vesicle proteome (Judith et al., 2019). Second, ATG9A is in INVs and ATG9 vesicles have an INV marker, as revealed by the reciprocal relocalization–co-relocation of ATG9A and TPD54 in cells. Third, some ATG9 vesicle cargos were also confirmed to be present in INVs using INV capture at the mitochondria. Fourth, depletion of TPD54 impaired the autophagy response: decreased LC3B lipidation, reduced autophagosome size and inhibition of subcellular redistribution of ATG9A. Fifth, comparison of vesicle capture efficiency suggests that approximately 20% of INVs are ATG9A-flavor, whereas all ATG9 vesicles that could be captured are INVs. Finally, ATG9A-flavor INVs could be redistributed selectively to the cell periphery by RUSC2 overexpression, whereas a CIMPR-flavor of INV was unaffected. These findings are supplemented by other evidence such as the similar diffusivity of ATG9A-containing vesicles and INVs (Sittewelle and Royle, 2024; Broadbent et al., 2023).

The established function of ATG9 vesicles is as a membrane reservoir for autophagosome formation during autophagy. Accordingly, we found that depletion of TPD54 impeded autophagosome formation and autophagic flux in starved cells. In addition, the loss of ATG9A from the TGN upon starvation was also inhibited by TPD54 depletion. This suggests that TPD54 may be involved in the formation of ATG9A-flavor INVs at the TGN. Previous work indicates that ATG9 vesicles form at the TGN by an AP-4-dependent mechanism (Davies et al., 2018; Guardia et al., 2021). Beyond a preference for high curvature membranes there has yet to be a function ascribed to TPD54, so it is unclear if it has a role in vesicle formation. A stringent analysis of proteins in dynamic organellar maps which are altered in response to loss of AP-4, identified TPD54 (Davies et al., 2022). This identification might simply reflect the effect of AP-4 knockout on ATG9A-flavor INVs, but it may also suggest that TPD54 works together with AP-4 in vesicle formation at the TGN. In support of this, the increase in LC3B lipidation upon depletion of TPD54 is reminiscent to that seen following knockout of AP-4 subunits AP4E1 and AP4B1 (Davies et al., 2018). On the other hand, an AP-4-specific function for TPD54 seems unlikely since depletion of TPD54 affects many other membrane trafficking steps – for example, recycling of integrins and transferrin receptor, anterograde traffic – which are not governed by AP-4 and would argue for a more ubiquitous role for this protein during vesicle formation (Larocque et al., 2020, 2021).

While ATG9 vesicles have been synonymous with autophagy, there has been a growing appreciation of their autophagy-independent functions. ATG9 vesicles have been shown to be involved in plasma membrane repair following exotoxin injury, and also in lipid mobilization from lipid droplets through association with the lipid transferase TMEM41B (Claude-Taupin et al., 2021; Mailler et al., 2021). In addition, Campisi et al. (2022) described a role for ATG9 vesicles in integrin trafficking during cell migration; a function which is remarkably similar to that attributed to INV-mediated integrin recycling in our previous work (Larocque et al., 2021). Rather than an autophagy-specific vesicle type, the classification of ATG9A-flavor INVs as a vesicle sub-type with lipid scramblase activity that can participate in a wide range of cell biological activities is probably a more accurate definition.

If ∼80% of INVs are not ATG9A-flavor, then what are the other flavors of INV? The INV proteome highlights a number of interesting cargos which might constitute new subtypes of INVs. For example, glucose transporter 1 (SLC2A1), is a typical cargo sorted by retromer (Steinberg et al., 2013). Vesicles containing such cargos may constitute novel INV flavors. Conversely, other small uncoated vesicles that have been previously been described may actually be flavors of INV. For example, synaptic-like microvesicles (SLMVs) are synaptic vesicle-sized secretory vesicles found in non-neuronal cells (Clift-O’Grady et al., 1990; Cameron et al., 1991; Régnier-Vigouroux et al., 1991). We found considerable overlap of the INV proteome with a SLMV dataset (Salazar et al., 2005), due to the presence of synaptic vesicle proteins and paralogs (SCAMP1, SCAMP2, SCAMP3, SYNGR2, SYPL1) in the INV proteome. An SLMV-flavor of INV would be in keeping with the observation of fusion of synaptophysin-containing INVs at the cell surface (Sittewelle and Royle, 2024). Intriguingly, classic work showed that expression of synaptophysin in non-neuronal cells causes the aggregation of SLMVs (Cameron et al., 1991). Recently, coexpression of synapsin and synaptophysin was shown to enhance the clustering of SLMVs (Park et al., 2021) and that ATG9 vesicles and SLMVs are each present in the synapsin clusters, yet remain separate (Park et al., 2023). Our work suggests that these vesicles – collectively – are INVs, and that this is strong evidence that separate INV flavors do exist and can remain segregated.

Exploring the variety of INV flavors and determining any interplay between them is an important goal in order to understand the roles of INVs in membrane trafficking in health and disease.

## Methods

### Molecular biology

The following plasmids were available from previous work: GFP-FKBP-TPD54, mCherry-FKBP-tagged TPD54 WT and R159E mutant, GFP-FKBP and mCherry-MitoTrap (Larocque et al., 2021); ATG9A-GFP (Sittewelle and Royle, 2024). HA-RUSC2 was a kind gift from Alex Davies (Davies et al., 2018).

For low expression of GFP-TPD54 WT or R159E in HeLa cells, a pEGFP-C1 plasmid with the CMV promoter exchanged for PGK was used, and a TPD54 cDNA was subcloned into this plasmid using XhoI and MfeI sites; and then the R159E mutation was introduced by site-directed mutagenesis. GFP-FKBP-TPD54 R159E was made by cloning TPD54 R159E into pEGFP-FKBP-C1 at XhoI and MfeI sites.

ATG9A-FKBP-mCherry was constructed by first making ATG9A-FKBP-GFP by amplification of human ATG9A and insertion at EcoRI and SalI sites in pFKBP-GFP-C1, followed by replacement of GFP with mCherry using NheI and NotI sites. Tagging ATG9A at the C-terminus was shown previously to be sufficient to functionally complement knockout of ATG9A (Mailler et al., 2021). FKBP-mCherry was made by replacing GFP in pFKBP-GFP-C1 with mCherry using NheI and NotI; FKBP-mCherry was cloned in place of mCherry using NheI/MfeI into a pmCherry-N1 vector with a crippled CMV promoter in order to reduce expression levels in cells. MitoTrap constructs were converted to the rapalog-specific FRB T2098L form by site directed mutagenesis of mCherry-MitoTrap and conversion to blue form by replacing mCherry with EBFP2 at AgeI/BrGI (Bayle et al., 2006). SH3GLB1-GFP, GFP-ARFIP2, DAGLB-GFP, GFP-PI4K2A, and GFP-PI4KB were each made by custom synthesis and cloning in pmEGFP-C1 or pmEGFP-N1 at EcoRI/SalI, EcoRI/KpnI, HindIII/SalI, EcoRI/SalI, and BglII/SalI sites respectively.

### Cell culture

Wild-type HeLa cells (HPA/ECACC 93021013) or GFP-TPD54 knock-in (clone 35) HeLa cells (Larocque et al., 2020) were maintained in DMEM with GlutaMAX and 25 mM HEPES (Thermo Fisher, 32430100) supplemented with 10% FBS, and 100 U mL^−1^ penicillin/streptomycin. All cells were kept in a humidified incubator at 37 °C and 5% CO_2_; and were routinely tested for mycoplasma contamination by PCR.

To generate cell lines stably expressing GFP-TPD54 WT or GFP-TPD54 R159E under a PGK promoter, HeLa cells were seeded at 50% confluency in a 75 cm^2^ flask and transfected with 5 µg of the respective plasmid, using Genejuice (Merck). After 48 h, media was changed to complete media supplemented with 500 µg mL^−1^ geneticin (Thermo Fisher, 10131035). Following selection, cells were assessed for GFP fluorescence by microscopy, expanded and were frozen down as stably transfected populations.

For transient transfection, 1.4 × 10^5^ cells were plated onto either coverslips or 35 mm glass bottom dishes (WPI – FD35-100). Plasmids were transfected using Genejuice and cells analyzed 36–48 h after transfection. For depletion of TPD54, siTPD54 (GUCCUACCUGU-UACGCAAU) or siGL2 (CGUACGCGGAAUACUUCGA) as a control were used (Larocque et al., 2020). Transfection of siRNA was by Lipofectamine RNAiMax according to the manufacturer’s instructions, with cells typically analyzed 48 h post-transfection.

### Isolation of intracellular nanovesicles

HeLa cells (14 × 10^6^) were seeded onto a 24.5 × 24.5 cm plate (Corning, 431110). Typically one plate per condition, per replicate was set up and left to grow until 80–100% confluent. For harvest, plates were placed on ice, cells were washed three times with 10 mL ice cold PBS and scraped into 5 mL PBS using a bisected rubber stopper. Cells were pelleted by centrifugation at 4200 *g* for 2 min at 4 °C and then lysed via resuspension of the cell pellet in 1 mL INV release buffer (20 mM Tris-Cl, 300 mM NaCl, 5 µg mL^−1^ digitonin, 0.2 mM PMSF, with cOmplete EDTA free proteinase inhibitor (Roche, 4693132001), pH 7.5) in a pre-cooled microcentrifuge tube. Cells were lysed on ice for 30 min, with gentle pipetting every 10 min using a 1 mL tip. During lysis, GFP-Trap Agarose beads (chromotek, gta-20), 25 µL per reaction, were prepared by washing three times with 500 mL INV release buffer with spins at 2500 *g* for 2 min at 4 °C before final resuspension in INV release buffer (50 µL per reaction). For experiments where a control IP was performed, Rho1D4 MagBeads (Genaxxon, S5394.0005) were used, and magnetic isolation performed. Lysates were then centrifuged at 20 000 *g* for 15 min at 4 °C, 50 µL supernatant was reserved for analysis and the remainder was incubated with GFP-Trap beads (50 µL per reaction) in a pre-cooled microcentrifuge tube and incubated with end-over-end rotation at 4 °C for 1 h. The bead-lysate mixture was centrifuged at 20 000 *g* for 2 min at 4 °C to pellet the beads. The supernatant was retained for analysis and the INVs on beads were washed three times with INV release buffer. Digitonin in the washes reduced background but potentially interfered with mass spectrometry so was omitted for some experiments. Captured material was eluted by resuspending beads in 50 µL 2× laemmli buffer (Alfa Aesar, J60015.AD) and heating to 95 °C for 5 min.

### Mass spectrometry

Samples were loaded onto a 4–15% precast polyacrylamide gel (Bio-Rad, #4561084) and run at constant voltage until all proteins had migrated into the resolving gel. The gel was stained with InstantBlue protein stain (Abcam, ab119211) for 1 h at room temperature (RT) with gentle agitation. Individual lanes were excised and diced (2–4 mm) and transferred to 1.5 mL microcentrifuge tubes per lane. Gel pieces were washed three times with 1 mL 50% ethanol (EtOH), 50 mM ammonium bicarbonate (ABC) for 20 min at RT with shaking at 650 rpm. The fragments were then dehydrated with 200 µL 100% EtOH for 5 min at RT. For efficient reduction of disulphide bridges and alkylation of cysteines, the gel pieces were next rehydrated in 100 µL 10 mM TCEP, 55 mM CAA in 50 mM ABC and incubated for 30 min at 70 °C with shaking at 650 rpm. Slices were washed a further three times in 1 mL 50% EtOH, 50 mM ABC for 20 min at RT with shaking at 650 rpm before dehydration with 200 µL 100% EtOH for 5 min at RT. Trypsin was made up to 2.5 ng µL^−1^ in 50 mM ABC and 80 µL was used to rehydrate the gel pieces for 10 min at RT. Buffer volume was increased by adding additional 50 mM ABC to ensure all pieces were submerged followed by digestion overnight at 37 °C. The following day, the solution around the gel containing the peptides was collected. Gel pieces were submerged in 25% ACN 5% formic acid solution and sonicated in a water bath for 10 min at RT. This process was performed three times with solution collection and pooling per sample. Finally, the solutions were filtered using a Costar Spin-X centrifuge tube filter (Sigma, CLS8169) to remove any residual gel and then peptides were concentrated in a speed-vacuum for 3 h at 45 °C and resuspended in 50 µL 2% acetonitrile, 0.1% trifluoracetic acid (TFA) before being transferred to mass spectrometry vials. Samples were stored at −20 °C until running on either a timsTOF or Orbitrap mass spectrometer (Proteomics Facility, University of Warwick).

LC-MS/MS raw data files were processed using MaxQuant (v2.0.1) (Max Planck Institute of Biochemistry). The peptide lists were searched against the reviewed UniProt human proteome database (retrieved May 2021) supplemented with the sequence of GFP-TPD54 and the MaxQuant common contaminant database. Peptide abundance was quantified using the MaxQuant Label-Free Quantification (LFQ) algorithm. Enzyme specificity for trypsin was selected with the allowance of up to two missed cleavages. All searches were performed with cysteine carbamidomethylation set as the fixed modification and oxidation of methionine and acetylation of the protein N-terminus set as the variable modifications. The initial precursor mass deviation was set as 20 ppm and the fragment mass deviation set as 20 ppm. For label-free quantification, we set a minimum ratio count of 2 with 3 minimum and 6 average comparisons.

### Cell treatments

To capture vesicles at the mitochondria, an inducible heterodimerization approach was used as described previously, with minor modifications (Larocque et al., 2020). Briefly, cells expressing FKBP-tagged proteins and FRB T2098L-MitoTrap constructs were treated with rapalog AP21967 (A/C Heterodimerizer, TaKaRa Bio, 635056 or 635057) at a final concentration of 5 µM in media. Rapamycin induces autophagy and, although this occurs on a much longer timescale than our experiments, rapalog was used in place of rapamycin for induced heterodimerization. For fixed cell experiments, rapalog application was via complete exchange of media and a time point of 5 min was used for fixation. For live cell imaging, cells were in Leibovitz L-15 media supplemented with 10% FBS immediately prior to imaging and rapalog application was done using a 1:5 addition of a 5X stock in L-15 media with FBS.

Autophagy was induced by 3 h starvation in HBSS (Sigma Aldrich, H6648) or EBSS (Thermo Fisher, 24010043). To assess autophagic flux, bafilomycin A1 (Sigma Aldrich, SML1661) was applied at a final concentration of 100 nM during this incubation. Control conditions were treated with a 1:1600 dilution of DMSO.

### Immunofluorescence

Cells on glass cover slips were fixed using 3% PFA/4% sucrose in PBS for 15 min. After fixation, cells were washed with PBS and then incubated in permeabilization buffer (0.1% v/v Triton X-100 in 1xPBS) for 10 min. Cells were washed twice with PBS, before 45–60 min blocking (3% BSA, 5% goat serum in 1xPBS). Antibody dilutions were prepared in blocking solution. After blocking, cells were incubated for 2 h with primary antibody, PBS washed (3 washes, 5 min each), 1 h secondary antibody incubation (not required for directly conjugated primaries), 5 min each PBS wash (3 washes), mounting with ProLong Gold or Vectashield vibrance. Primary antibodies used: anti-ATG9A [EPR2450(2)], rabbit (Abcam, ab108338 or ab206253); anti-LC3B [EPR18709], rabbit (Abcam, ab192890); anti-CIMPR [2G11], mouse (Abcam, ab2733); anti-TGN46, sheep (Bio-RAD, AHP500G); anti HA [6E2], mouse (Cell Signaling, 2367); anti-HA [C29F4], rabbit (Cell Signaling, 3724); anti-GFP, rabbit, conjugated to Alexa Fluor488 (Thermo Fisher, A-21311). Secondary antibodies were Alexa Fluor568 or 647 conjugated goat anti-rabbit or anti-mouse, and donkey anti-sheep Alexa Fluor568, all highly cross-adsorbed (Thermo-Fisher). Note, for anti-LC3B staining, ice-cold methanol was used as a fixative (10 min) with the permeabilization step omitted.

### Western blotting

For Western blotting, cells were washed with ice-cold PBS and pelleted at 300 *g* for 5 min at 4 °C. Lysates were prepared using RIPA buffer (Thermo Fisher, 89900) supplemented with cOmplete EDTA-free protease inhibitor cocktail tablet (Roche, 11836170001), 0.2 mM PMSF and DNAse I (New England Biolabs, M0303L) 150 µL per 10 mL; for 30 min at 4 °C with gentle agitation. Protein concentrations were determined using the BCA assay, and samples were heated at 65 °C or 95 °C in Laemmli buffer for 5 min, and resolved on a precast 4–15% polyacrylamide gel (Bio-Rad). Proteins were transferred to nitrocellulose or PVDF using a iBlot2 Dry Blotting System (Bio-Rad). Following blocking in 5% w/v non-fat milk (Merck, 70166) in TBST buffer (20 mM Tris, 150 mM NaCl, 0.1% v/v Tween-20, pH 7.6), membranes were incubated with primary antibodies: rat monoclonal anti-GFP (Proteintech, 3H9), rabbit polyclonal anti-vinculin (Sigma Aldrich, V4139), rabbit polyclonal anti-LC3B (Merck, L7543) or rabbit polyclonal anti-TPD54 (Dundee Cell Products) at 1:1000 in 2% Milk TBST for 2 h at RT or at 4 °C overnight with agitation. After three washes in TBST, secondary antibodies, HRP-conjugated goat anti-rat IgG (Sigma Aldrich, A9037) or mouse anti-rabbit IgG (Santa Cruz Biotechnology, sc-2357) at 1:5000 in 2% milk TBST were applied for 1 h at RT with agitation. Blots were imaged using Amersham ECL Prime Western Blotting Detection Reagent (Cytiva, RPN2236) on a ChemiDoc MP (Bio-RAD) digital imaging system.

### Microscopy

All images were captured using a Nikon CSU-W1 spinning disc confocal system with SoRa upgrade (Yokogawa) with a Nikon, 100 *×*, 1.49 NA, oil, CFI SR HP Apo TIRF with optional 2.8 *×* intermediate magnification and a 95B Prime camera (Photometrics). The system has a CSU-W1 (Yokogawa) spinning disk unit with 50 µm and SoRa disks (SoRa disk used), Nikon Perfect Focus autofocus, Okolab microscope incubator, Nikon motorized xy stage and Nikon 200 µm z-piezo. Excitation was via 405 nm, 488 nm, 561 nm and 638 nm lasers with 405/488/561/640 nm dichroic and Blue, 446/60; Green, 525/50; Red, 600/52; FRed, 708/75 emission filters. Acquisition and image capture was via NiS Elements software (Nikon). All microscopy data was stored via automated nightly upload to an OMERO database in the native file format (nd2).

### Data analysis

Data from MaxQuant (proteinGroups.txt files) were processed using VolcanoPlot in Igor Pro 9 (Royle, 2024). Processing in this package mimics the workflow in Perseus however, it allows multiple proteinGroups files to be easily combined. For the WT vs R159E vs Control INV comparisons, two files of three replicates of three conditions each were processed together. For the GFP-TPD54 knock-in INV isolation, three files of three replicates of two conditions each where the GFP-Trap isolation was compared to a Rho1D4-MagBead control isolation, were processed together. We used a cut-off of two-fold enrichment and *p <* 0.05 to determine INV proteins.

The outputs from VolcanoPlot were used for further analysis in R. Briefly, the two INV proteomes and their respective backgrounds were consolidated. For each protein, the subcellular location of “secreted” and the count of transmembrane domains in each group was retrieved from Uniprot. To classify INV proteins, PANTHER 18.0 protein classifications were retrieved using Bioconductor/PANTHER.db (Muller, 2017). The two topmost hierarchies of classification (below protein class) were used to construct the treemap. Many characterized proteins have no PANTHER classification, and so some “unclassified” proteins were manually assigned. Enriched analysis of Gene Ontology (GO) terms was done using Bioconductor package clusterProfiler (Wu et al., 2021).

For comparison with SLMV, ATG9 vesicle and CCV datasets, we used published data and read the data into R (Judith et al., 2019; Salazar et al., 2005; Borner et al., 2012). For SLMV data, we mapped all rat proteins to their human counterparts; for ATG9 data, we used 0.5 (log2) difference to remove background; and for all datasets we used Gene Name to match co-occurrence in datasets.

For relocalization–co-relocation analysis, three-channel images (FKBP protein, POI, Mitotrap) were first registered with NanoJ using TetraSpeck bead images (Laine et al., 2019). The mitochondrial channel was segmented using a Labkit classifier trained on MitoTrap images (Arzt et al., 2022). A ‘cytoplasm’ mask was then made by excluding the mitochondrial mask from a duplicate mask that had been dilated eight times. Measures of the FKBP and POI channels were taken in these two regions and background-subtracted to give the fluorescence of those two regions (F_mito_ and F_cyto_). Relocalization and co-relocation are shown in two ways. First, for live cell experiments, the fluorescence in the mitochondrial region post-treatment is divided by the pre-treatment value (F_post_/F_pre_), for each cell. Second, for fixed cell experiments where this is not possible, rapalog-treated cells are compared to control cells using the fluorescence at the mitochondria divided by the sum of the fluorescence in mitochondrial and cytoplasmic regions (F_mito_/F_total_). In each case, any negative responding cells (F_mito_/F_cyto_ < 1, FKBP channel, post-treatment) were removed from the analysis. In order to compare cytoplasmic loss of fluorescence due to relocalization– co-relocation, F_cyto_/F_total_ was calculated from fixed cell or live cell data using rapalog-treated cells. Where dynamic live cell data was quantified, the mitochondrial fluorescence was read in IgorPro and used for curve fitting using equation 1.

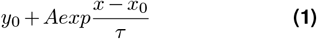

Traces are presented as the mitochondrial fluorescence (F) divided by the fluorescence at the beginning of the recording (F_0_), as F/F_0_.

For LC3 puncta quantification, normalized image stacks were segmented in Fiji using the Labkit plugin. The stacks were scaled isotropically, the segmented puncta were labeled using ‘connected components’ and their statistics retrieved using CLIJ (Haase et al., 2020). LC3 puncta were defined as >0.012 µm^3^. To quantify ATG9A at the Golgi, the two fluorescence channels were first registered using NanoJ using TetraSpeck bead images (Laine et al., 2019). The TGN46 channel was normalized and segmented in Fiji using a Labkit classifier trained on a subset of images. The resulting masks were used to quantify the mean pixel density in the ATG9A channel. Unless stated otherwise, image analysis outputs from ImageJ were read into R, analyzed and plotted using custom-written scripts.

## Data and software availability

All code used in the manuscript is available at https://github.com/quantixed/p064p037.

## ACKNOWLEDGEMENTS

We thank Sean Munro and Jerôme Cattin-Ortolá for sharing unpublished data with us, and Sharon Tooze for sharing reagents and helpful discussions. We are indebted to Cleidi Zampronio in the Proteomics Facility RTP and to the Computing and Advanced Microscopy Unit (CAMDU) for their help and support. We are grateful to all members of the Royle lab for feedback and critical discussion. The work was supported by grants from UKRI-BBSRC (BB/V003062/1) and Human Frontier Science Program (HFSP RGP25/2022). MF and DJM were supported by UKRI-MRC Studentships (MR/N014294/1). We thank Julius Muller for updating PANTHER.db to work with the latest PANTHER version. For the purpose of open access, the authors have applied a Creative Commons Attribution (CC BY) license to any Author Accepted Manuscript version arising

## AUTHOR CONTRIBUTIONS

MF, all ATG9 cell biological experiments, data analysis, programming; DJM, developed INV isolation and proteomic methods; PE, deep INV proteomic dataset; EC, LC3 western blotting; SJR, data analysis, programming, wrote the manuscript.

## Supplementary Information

**Figure S1.**
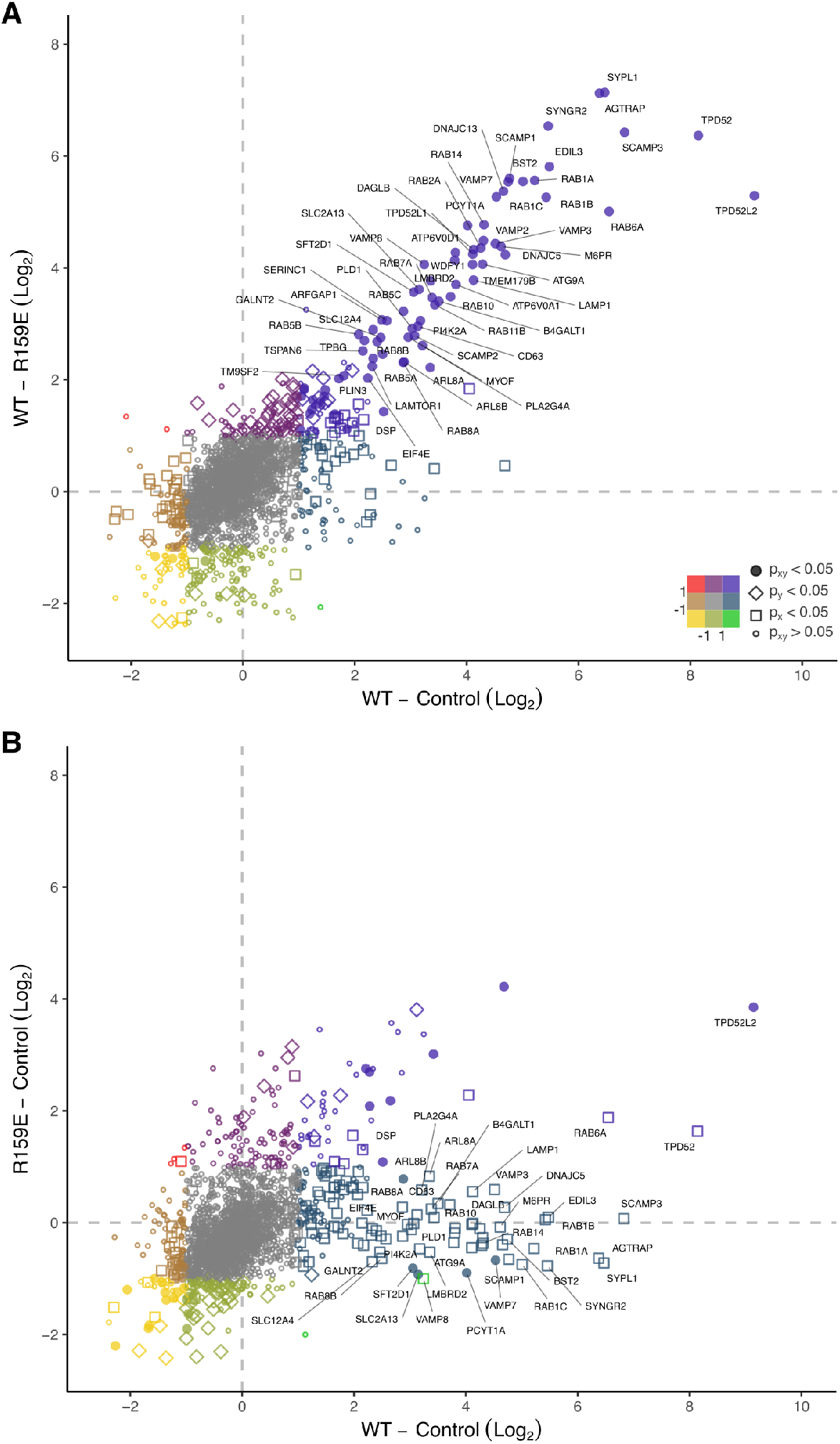
Verifying the INV proteome. (**A**,**B**) INV protein enrichment data. Data from Figure 1 replotted to compare enrichment in WT vs R159E against WT vs Control pulldown (A), or R159E vs Control against WT vs Control pulldown (B). Proteins significantly enriched in WT vs Control are labeled sparsely. Colors indicate the fold-enrichment (log2) while shapes indicate p-values of the comparison, as indicated.

**Figure S2.**
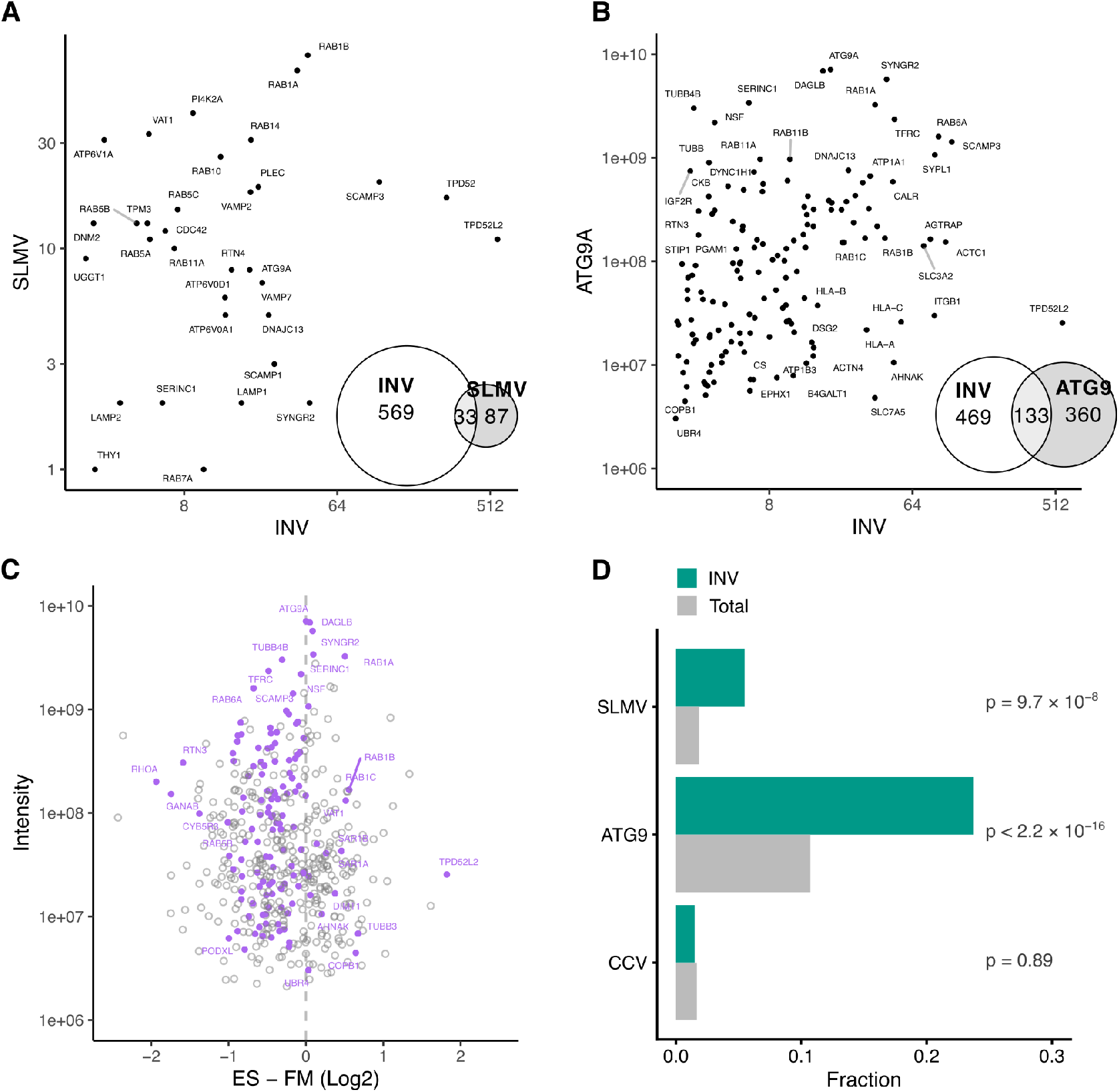
Comparison of the INV proteome with published vesicle datasets. (**A**) Comparison of INV enrichment data with SLMV proteomic dataset (Salazar et al., 2005). Log-log plot of SMLV peptide values vs INV enrichment data. (**B**) Comparison of INV enrichment data with ATG9 proteomic dataset (Judith et al., 2019). Log-log plot of ATG9 intensity values vs INV enrichment data. Insets: Euler plot to show the the number of proteins in each dataset and their overlap. (**C**) Proteins associated with ATG9A-positive membranes from cells incubated in full media (FM) or EBSS for amino acid depletion (ES). Proteins that are enriched in the INV dataset are highlighted purple. (**D**) Fractions of INV proteins and of total proteins that were featured in the indicated dataset. Enrichment of dataset proteins in INVs versus total is indicated, p-values from Pearson’s Chi-squared test.

**Figure S3.**
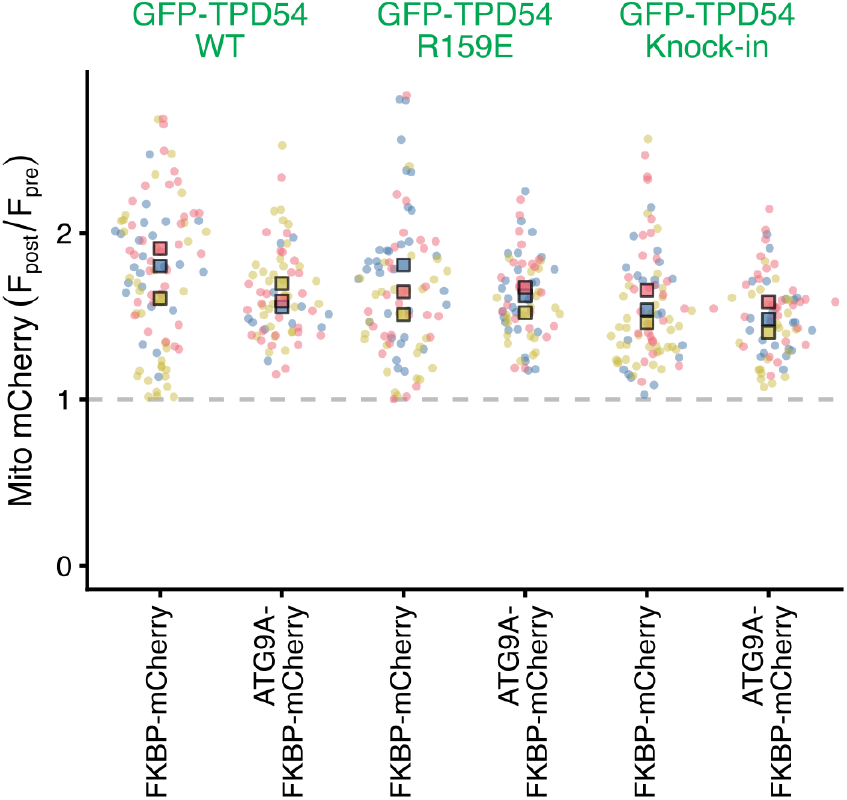
Relocalization of FKBP-mCherry or ATG9A-FKBP-mCherry. Superplot to show the ratio of mitochondrial fluorescence of FKBP-mCherry or ATG9A-FKBP-mCherry post- vs pre-rapalog treatment. Spots indicate individual cell measurements (n = 21 - 69 per repeat), colors indicate independent experimental repeats (n = 3), squares show the mean value for each replicate. Co-relocation results for the green channel are shown in Figure 4.

**Figure S4.**
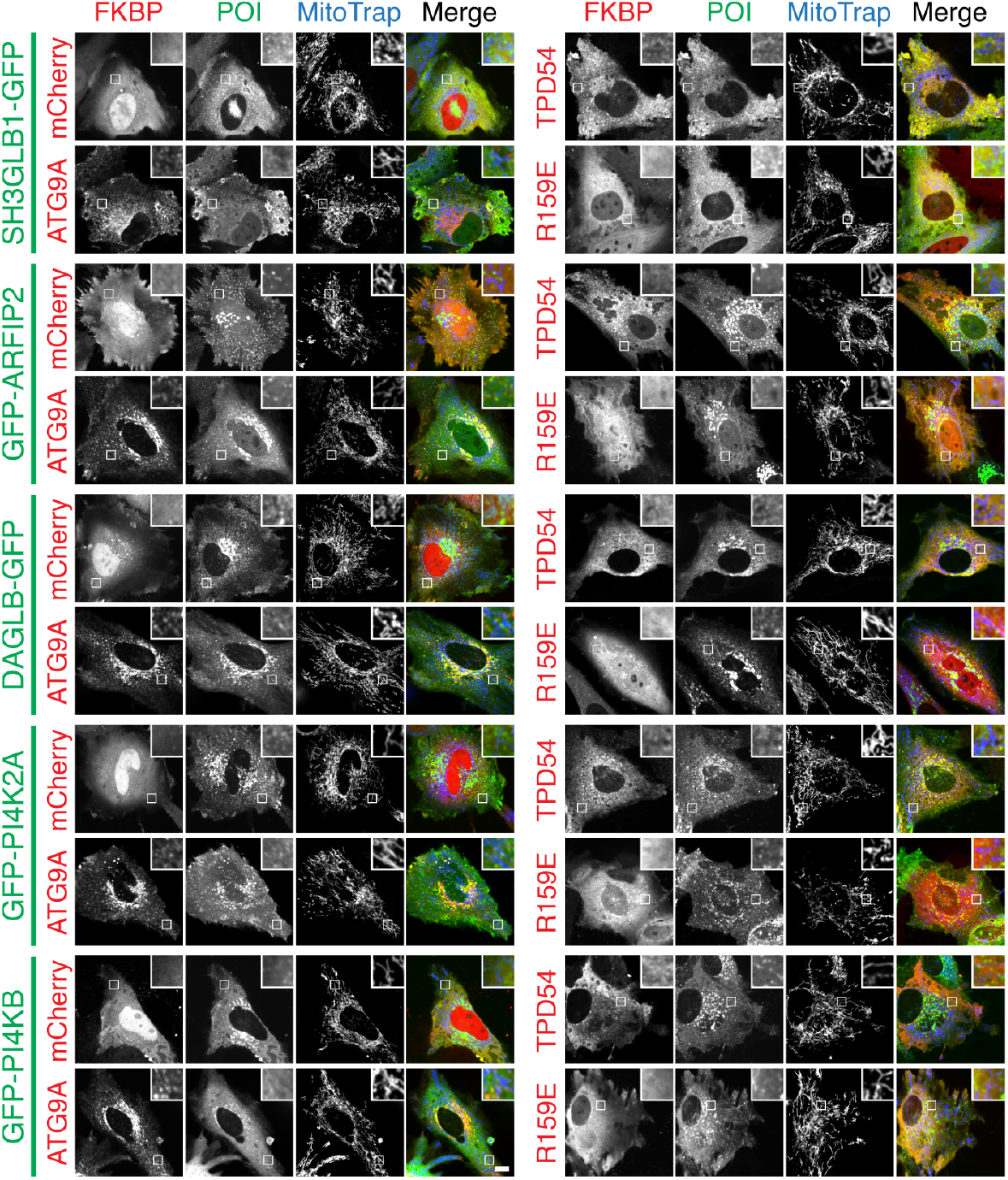
ATG9A-flavor INVs have ATG9 vesicle cargos – pre-rapalog images. (**A**) Representative confocal images of HeLa cells expressing MitoTrap (pMito-EBFP2-FRB T2098L, blue) and either FKBP-mCherry, ATG9A-FKBP-mCherry, mCherry-FKBP-TPD54 WT, or mCherry-FKBP-TPD54 R159E (red). Cells are co-expressing GFP-tagged SH3GLB1, ARFIP2, DAGLB, PI4K2A, or PI4KB as indicated (green). For each condition, only the pre-treatment image is shown, for the post-rapalog images see Figure 6. Scale bar, 10 µm; insets, 4 *×* zoom.

**Figure S5.**
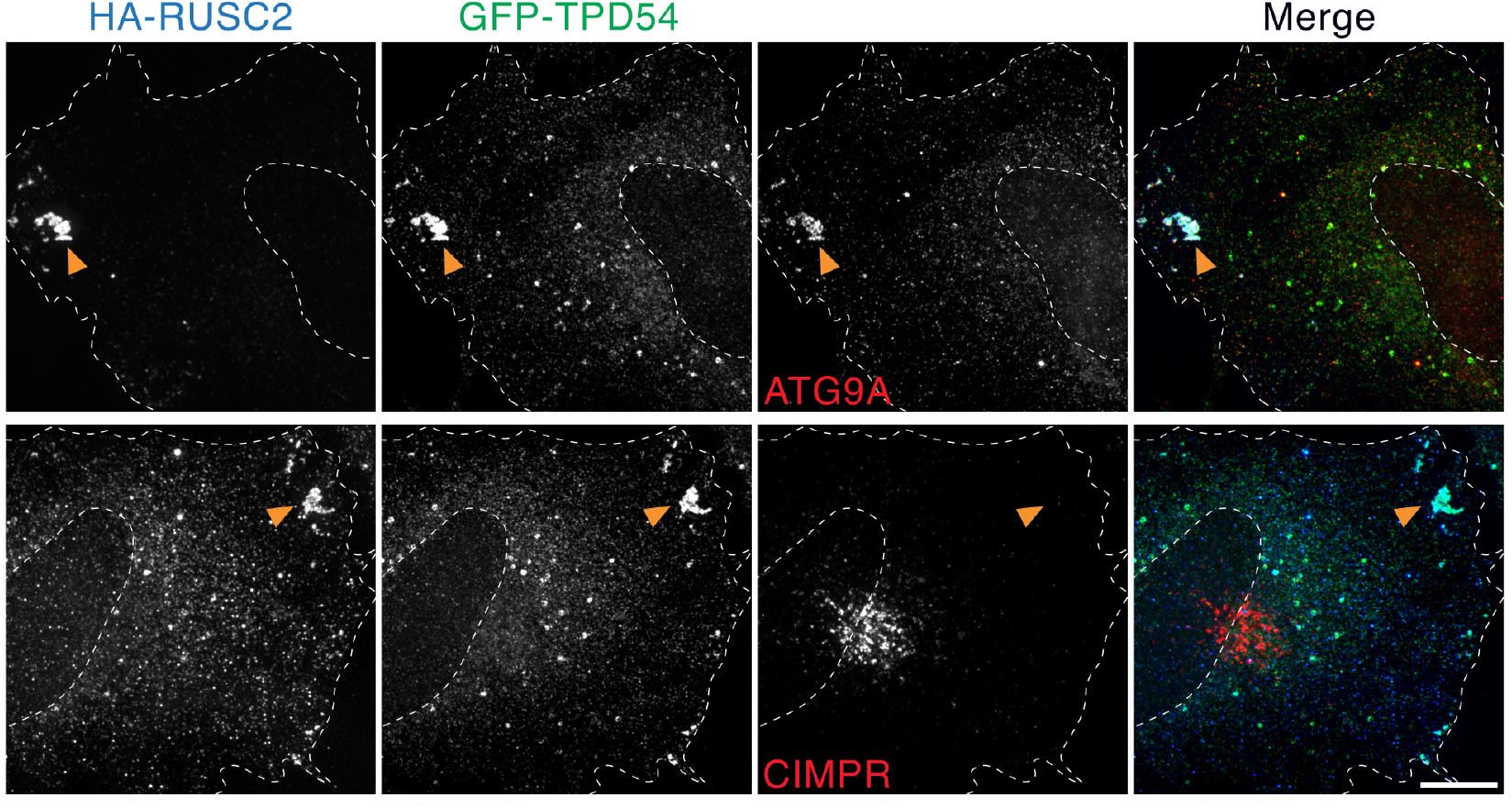
Selective relocalization of ATG9A-flavor INVs using RUSC2. Representative confocal micrographs of GFP-TPD54 (green) knock-in HeLa cells overexpressing HA-RUSC2 (blue). Cells were fixed and stained for ATG9A or CIMPR (red, as indicated), anti-HA as well as GFP boost. Orange arrowheads indicate accumulation of vesicular material at the cell periphery. Scale bar, 10 µm.

**Figure S6.**
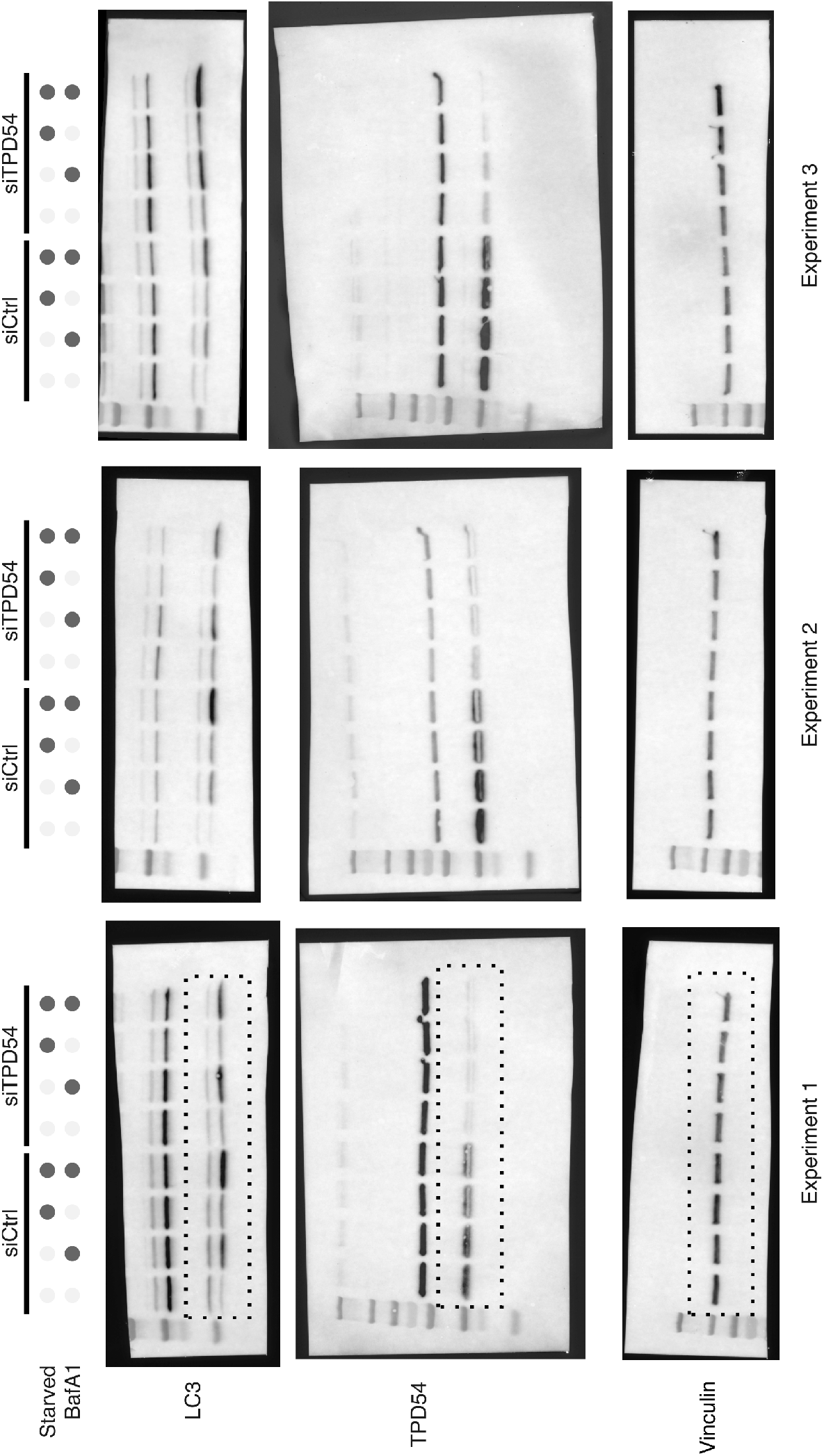
Effect of TPD54 depletion on LC3 lipidation under starvation. Three experiments where cells were transfected with siCtrl or siTPD54, starved (3 h) or not (fed) and either treated with BafA1 (100 µM) or not (DMSO), as indicated. LC3 was detected with anti-LC3B antibody, endogenous TPD54 was detected using an anti-TPD54 antibody; detection of Vinculin was used as a loading control. Dotted lines indicate the crops shown in Figure 7.

**Figure S7.**
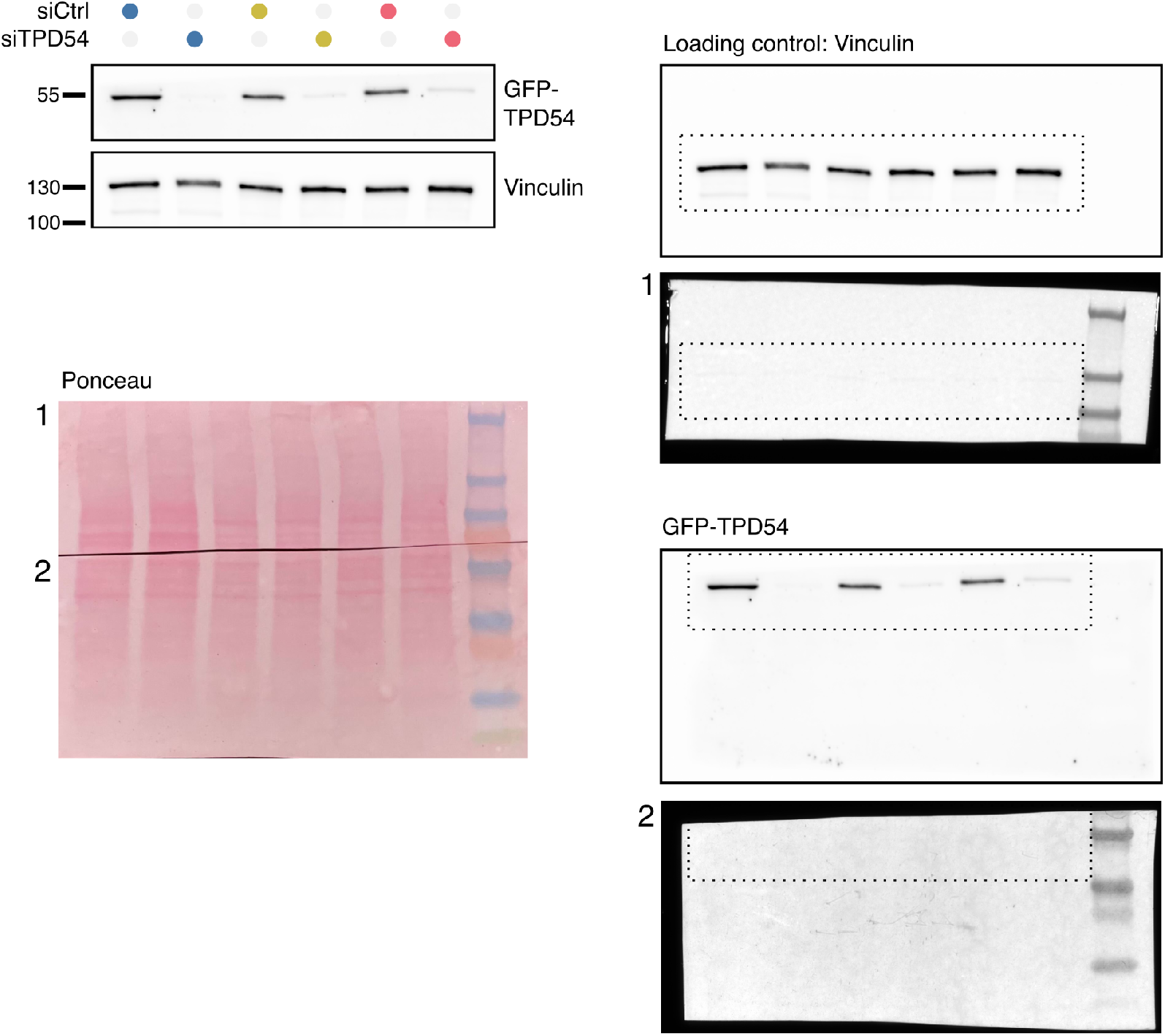
Depletion of TPD54 using RNAi. Western blot to show knock-down of TPD54 in the experiments shown in Figures 7 and 8). Endogenous GFP-TPD54 was detected using an anti-GFP antibody and vinculin was used as a loading control. Markers in kDa, colors indicate experimental repeats in Figures 7 and 8) and siRNA treatment. Other panels show the full Ponceau stained membrane and the full imaged area.

## Supplementary Videos

**Figure SV1.**
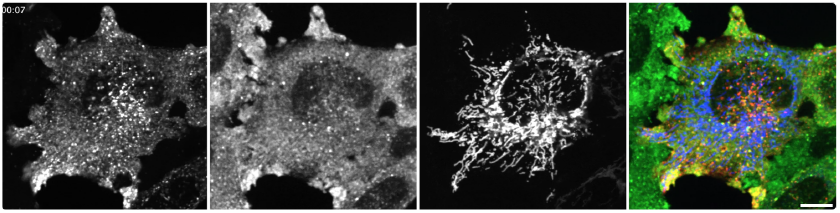
Reclocalization of ATG9A-FKBP-mCherry causes co-relocalization of endogenous GFP-TPD54. Movie of GFP-TPD54 (panel 2, green) knock-in cells expressing ATG9A-FKBP-mCherry (panel 1, red) and MitoTrap (panel 3, blue); capture of ATG9A-positive vesicles at the mitochondria is induced by rapalog (5 µM at 40 s). Time, mm:ss. Playback, 10 fps. Scale bar, 10 µm.

## Supplementary Tables

**Table S1.**
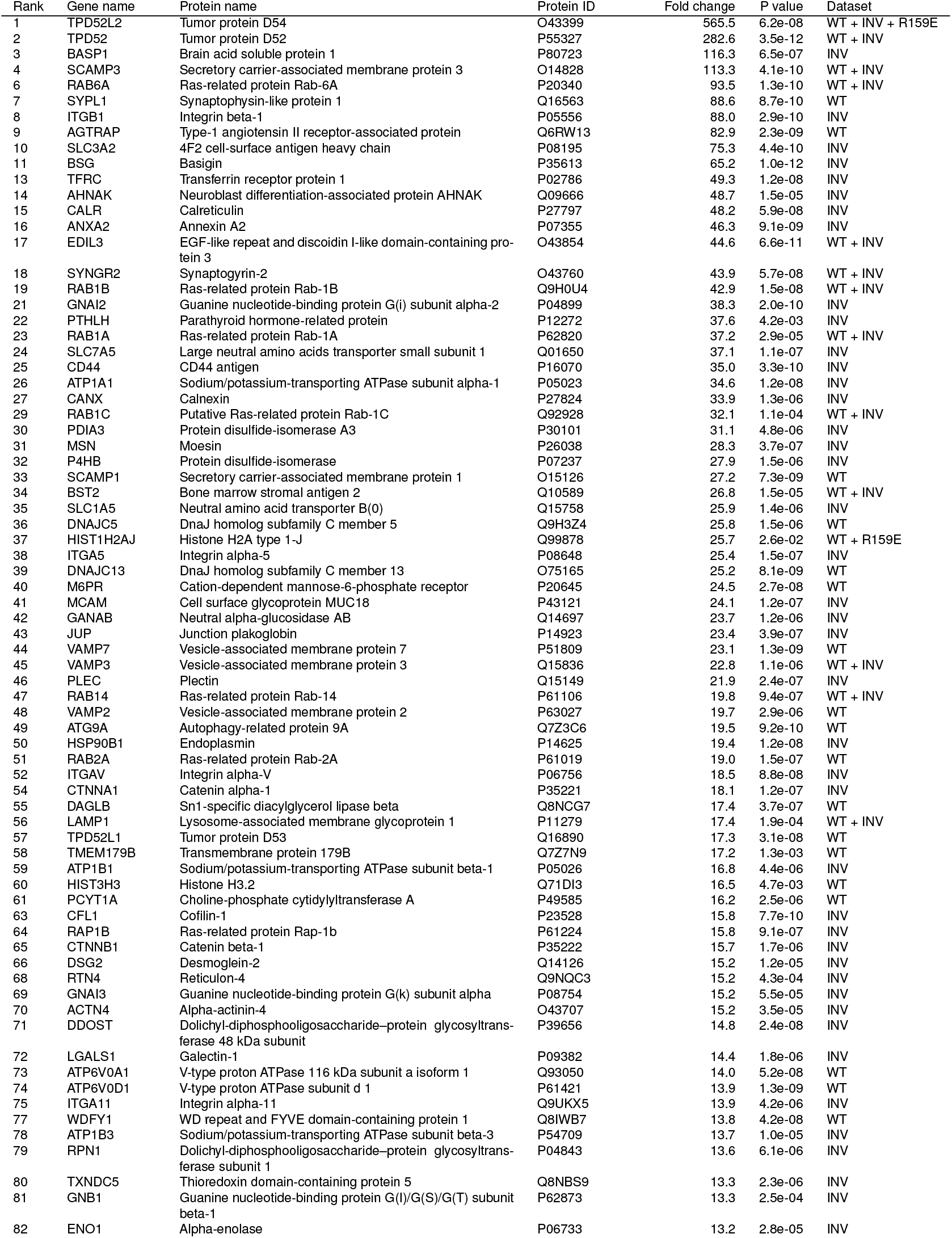
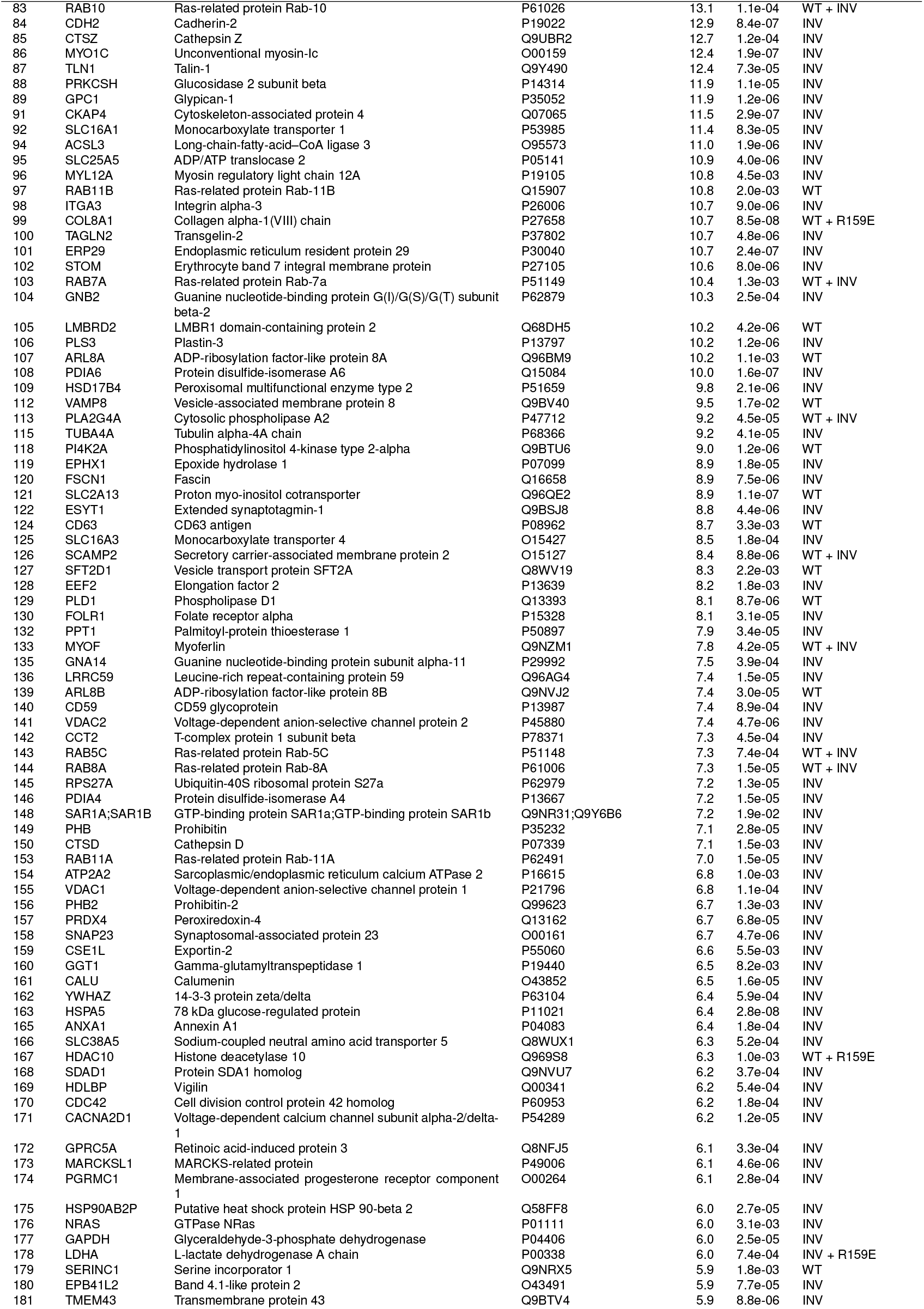
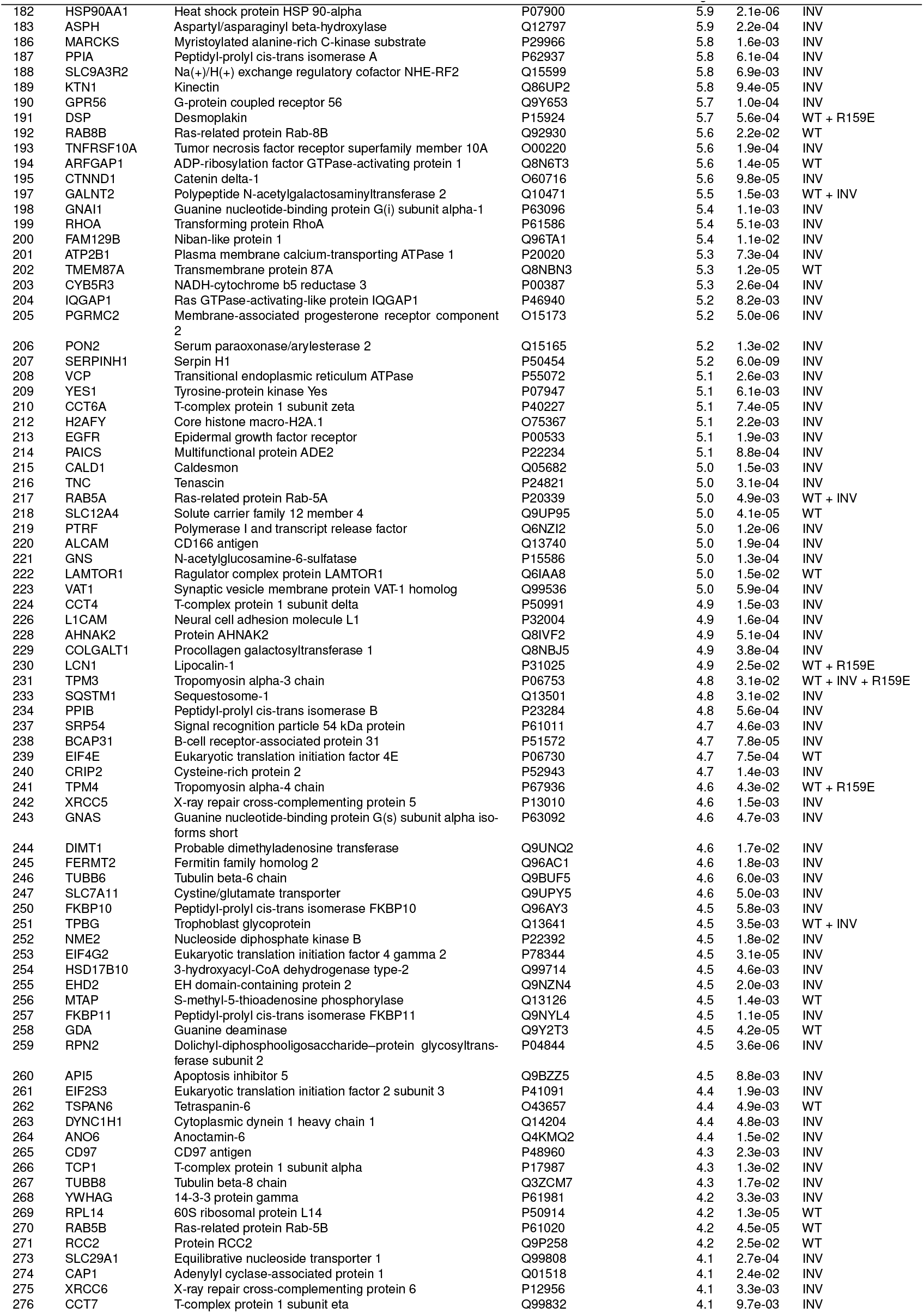
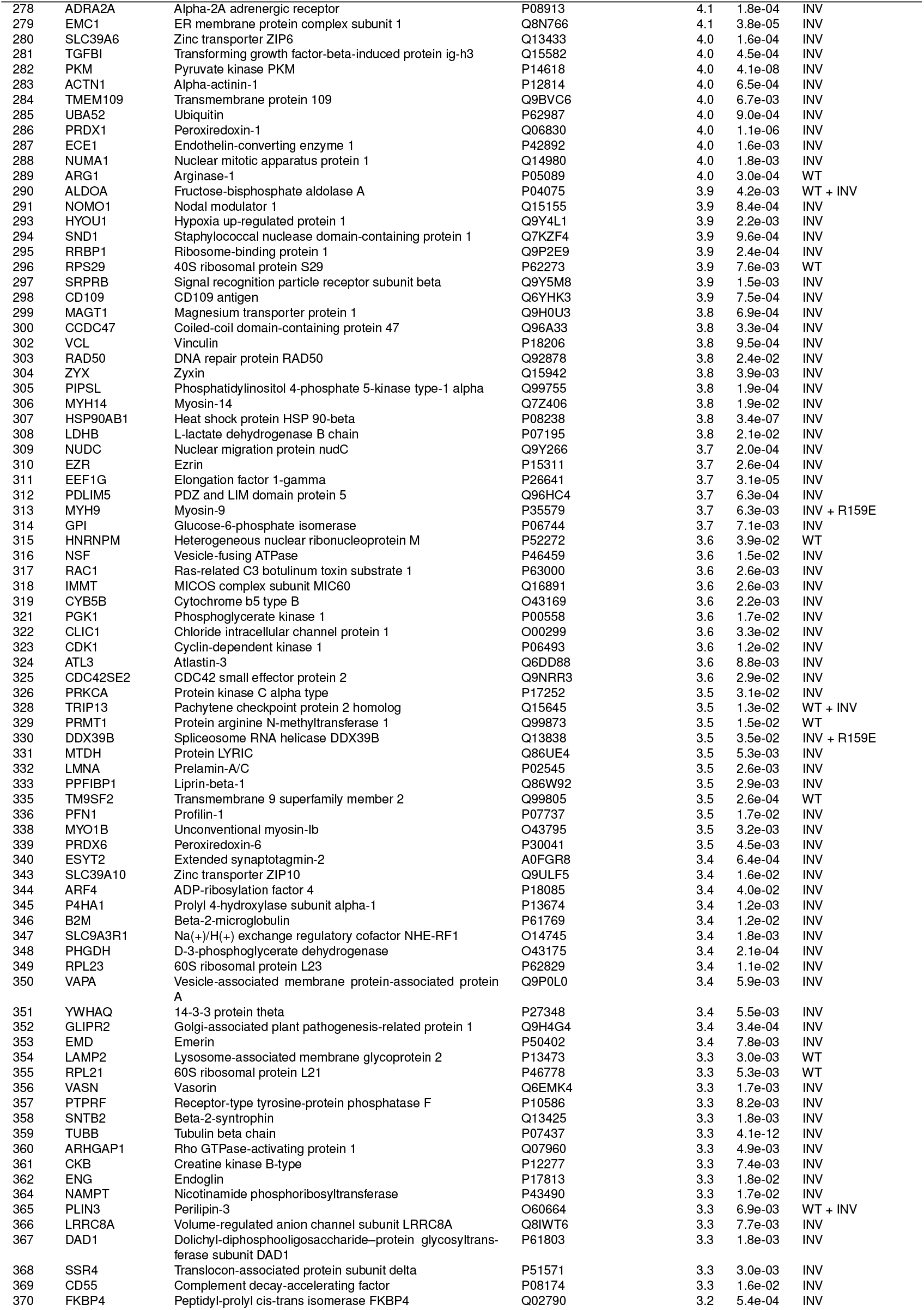
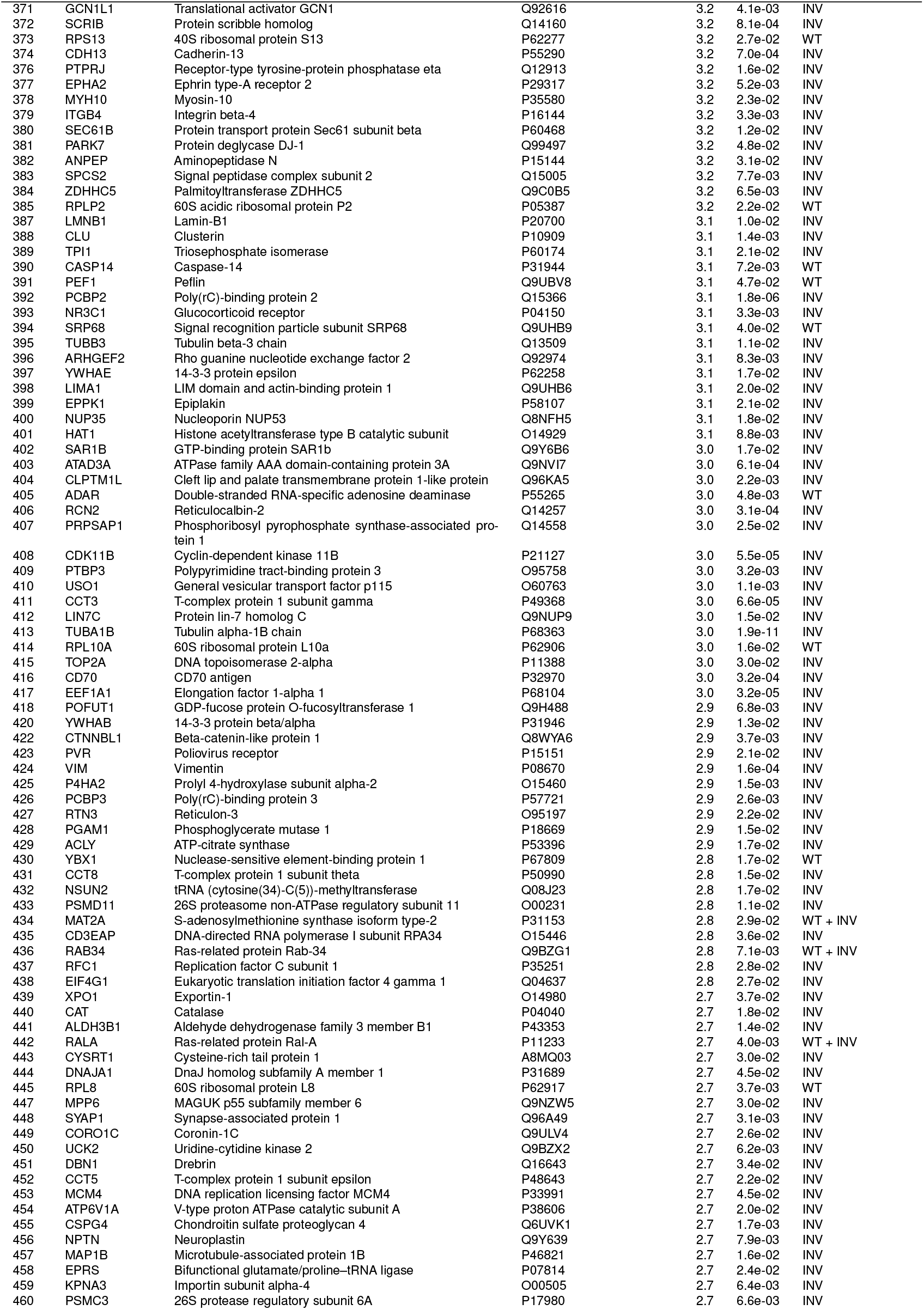
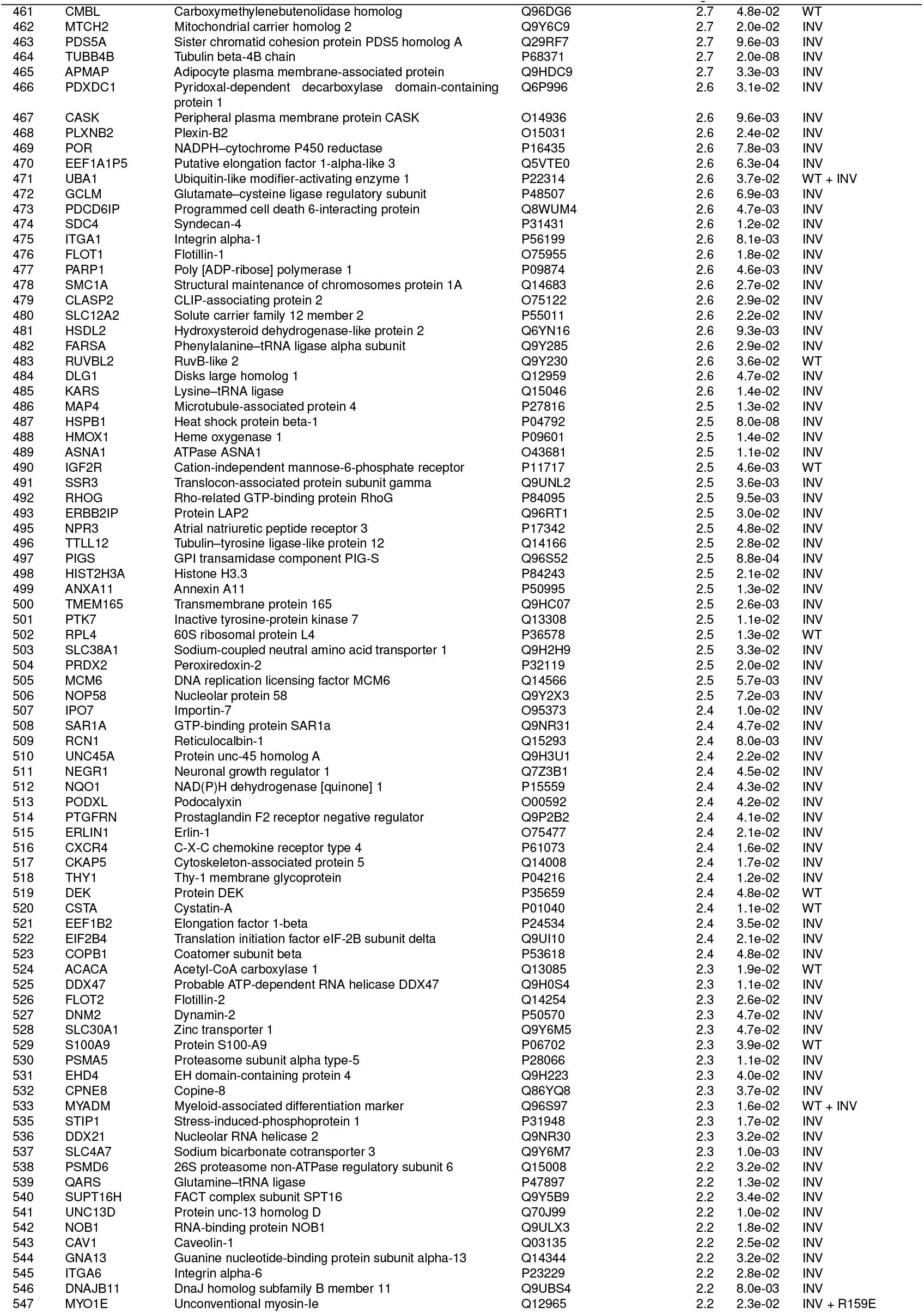
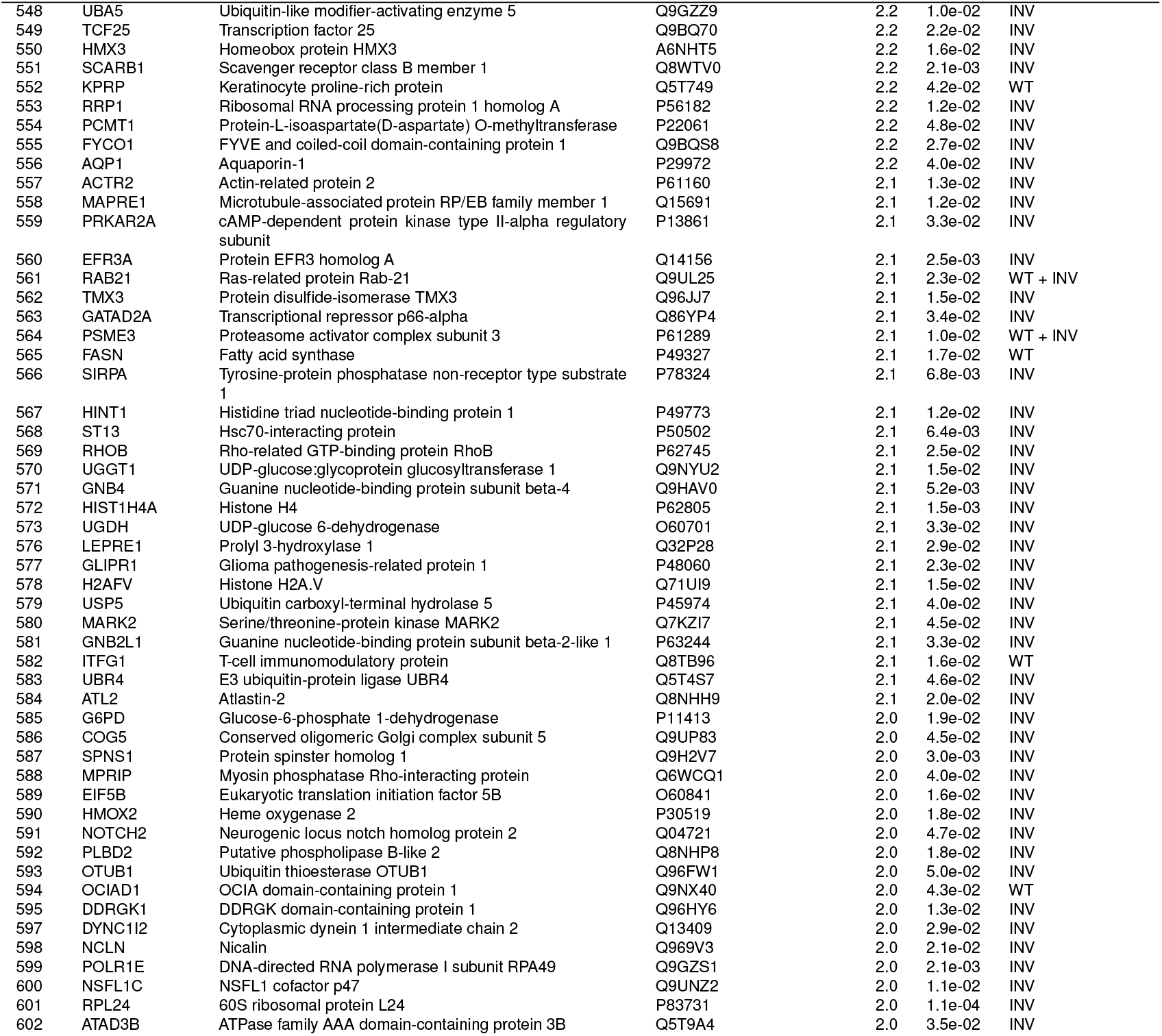
The INV proteome. A consolidated list of INV proteins determined by proteomics ranked by their fold enrichment over control. 602 proteins that had a fold change of > 2 and p < 0.05 are included. The enrichment in WT, R159E and/or knock-in over their respective controls is indicated as WT, R159E, and/or INV, respectively.

**Table S2.**
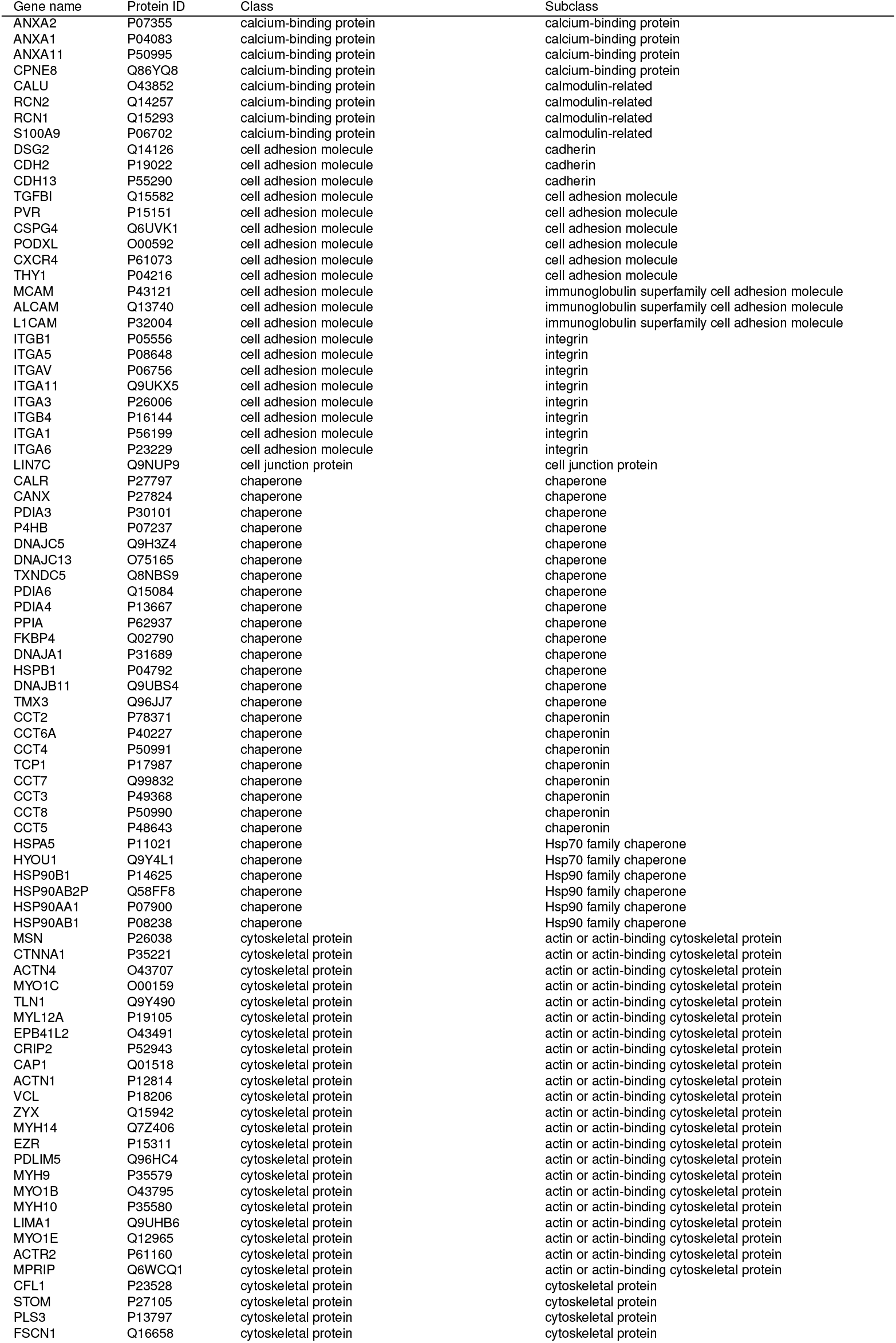
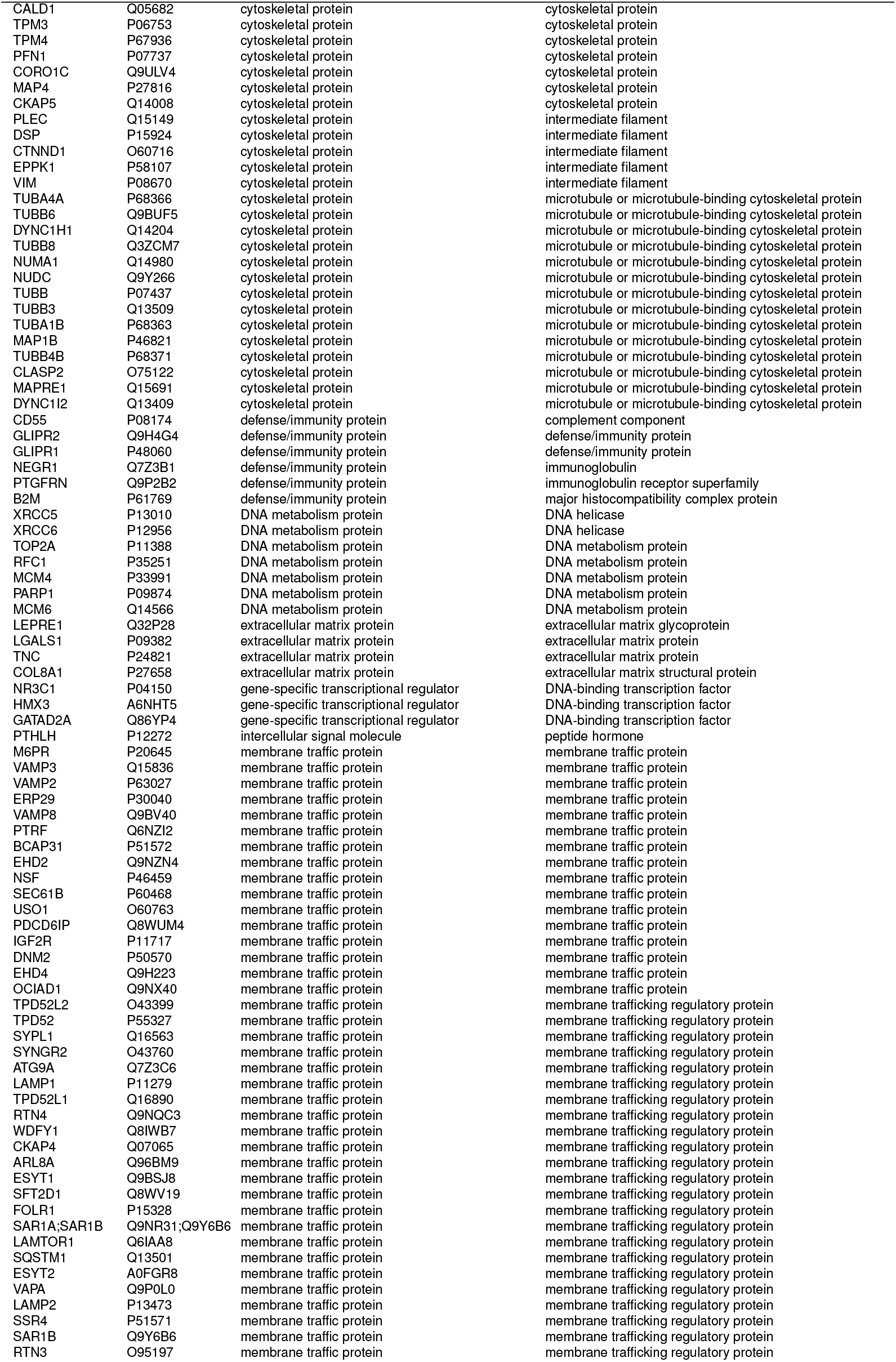
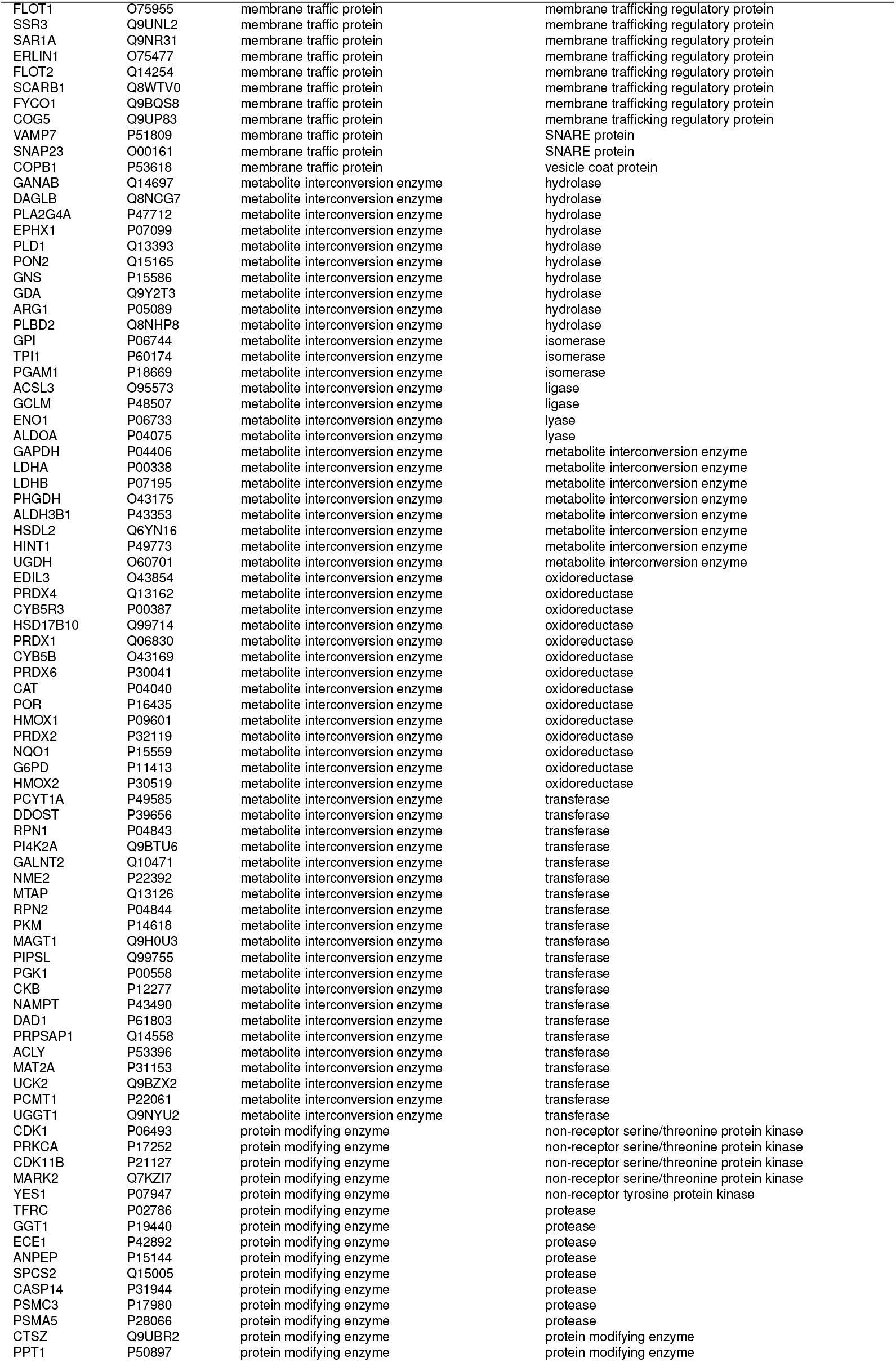
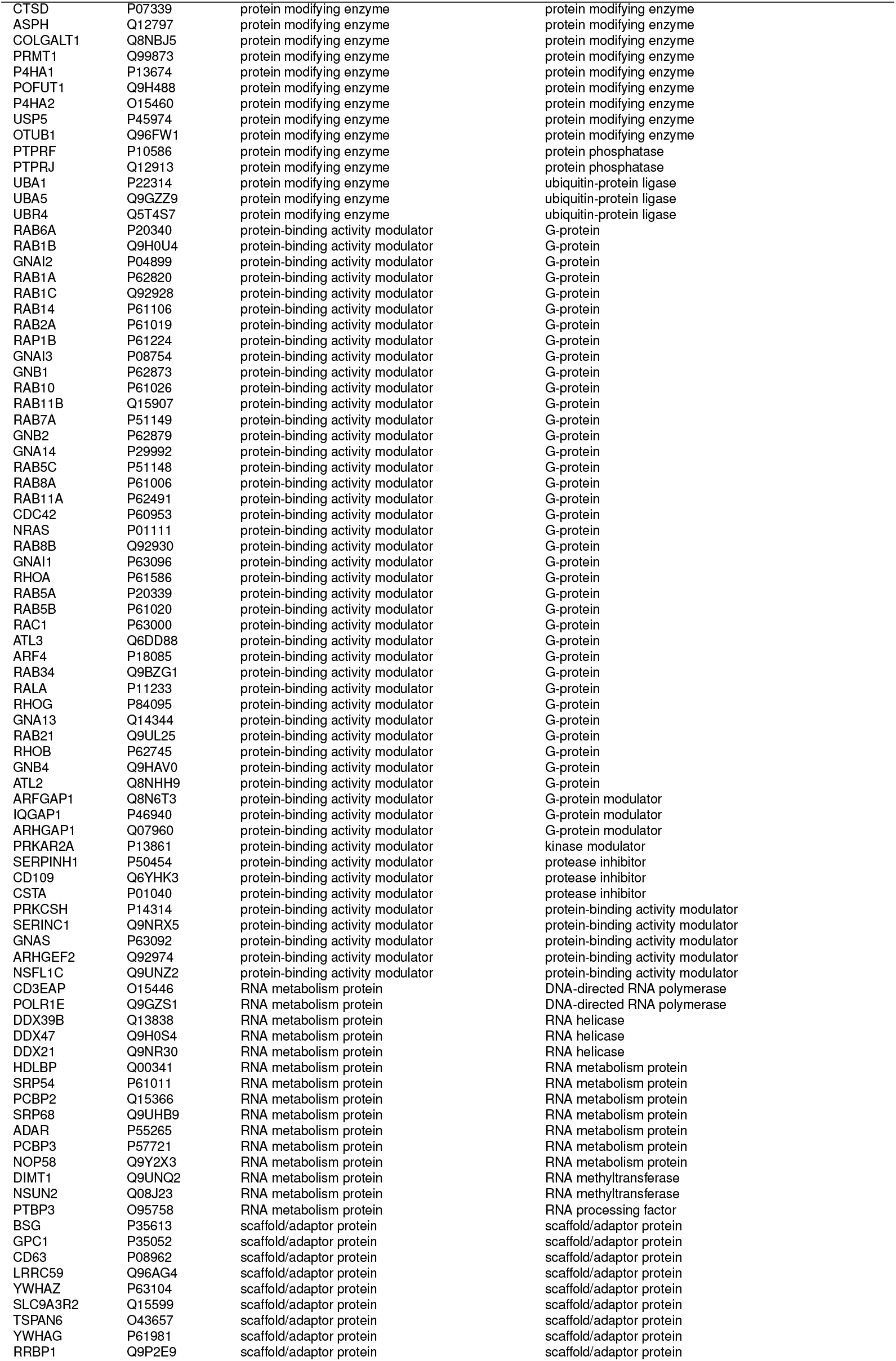
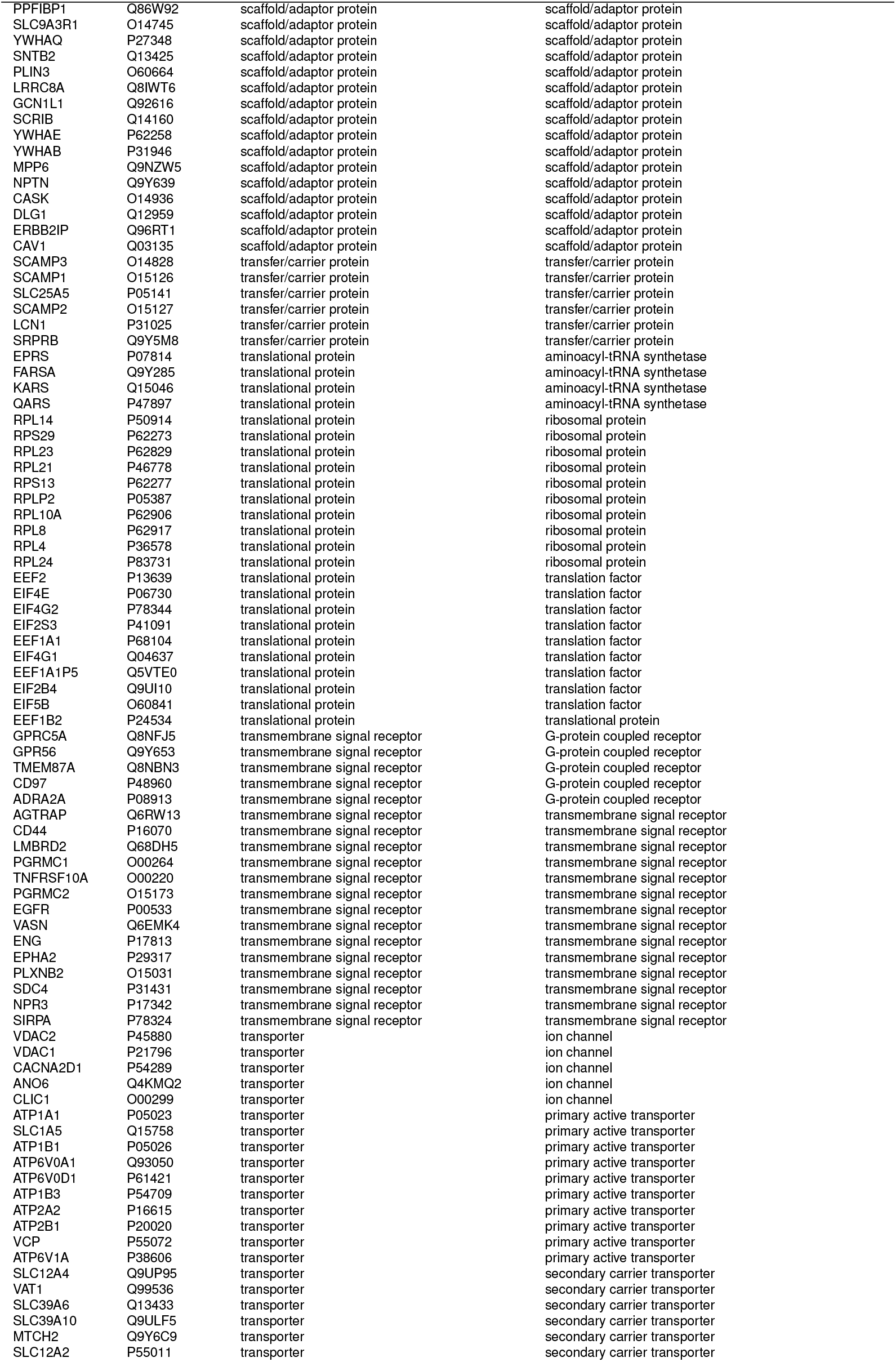
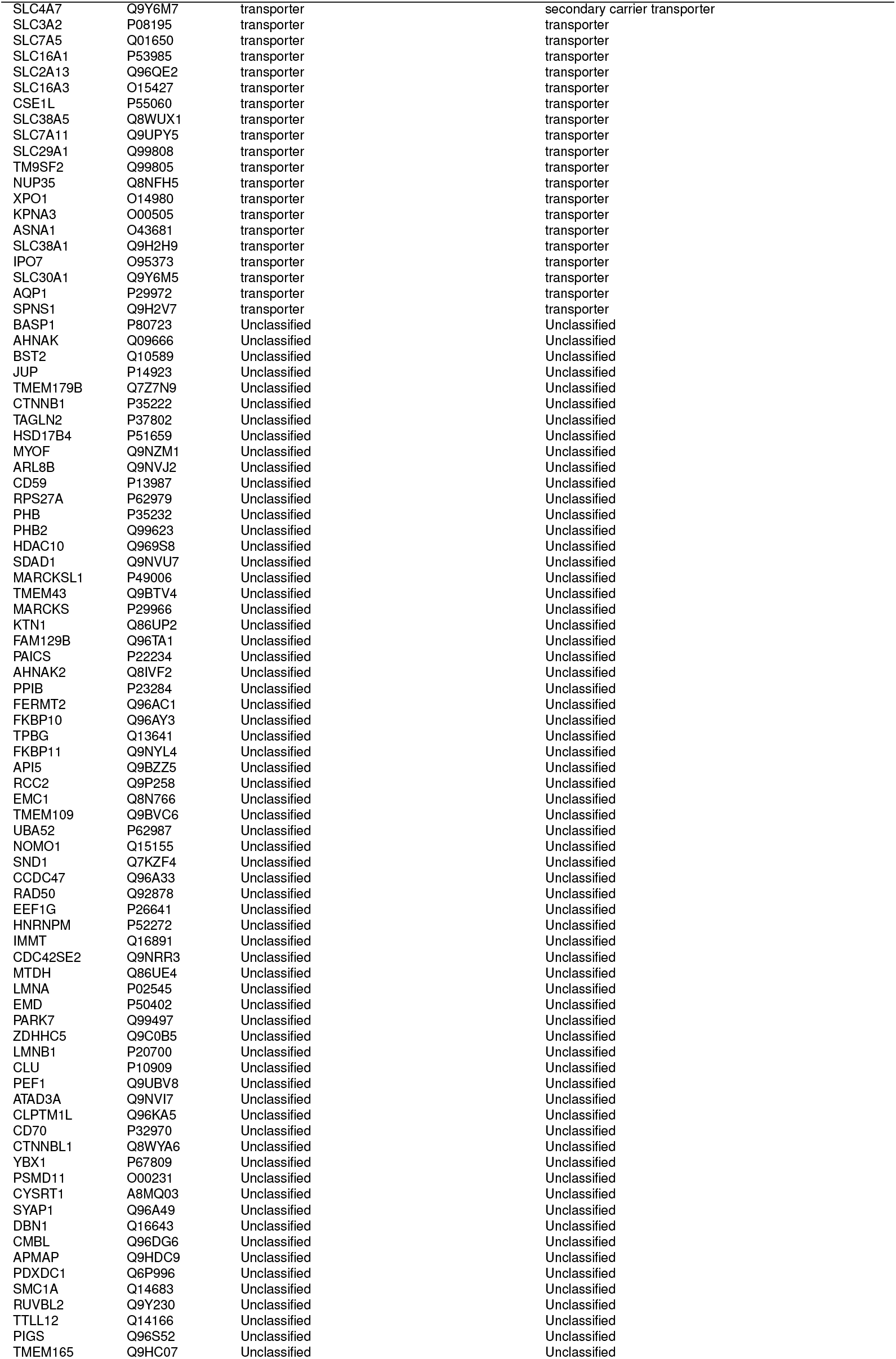
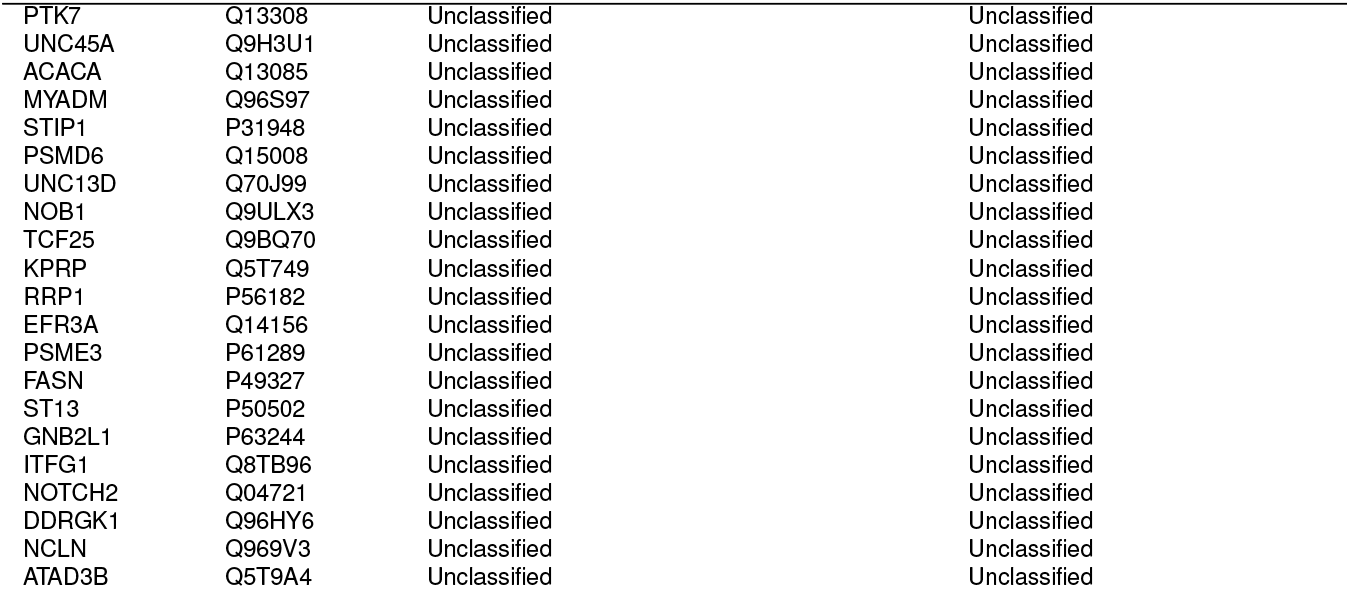
PANTHER protein classification of INV proteins. The consolidated INV proteome with assigned class and subclass from PANTHER protein classification are shown ordered by class-subclass.

